# A single-nucleus multiomic and spatial atlas of gene regulation in human spermatogenesis

**DOI:** 10.64898/2026.09.04.749409

**Authors:** Jahnavi Bhaskaran, Elizabeth Ing-Simmons, Irina Balaguer Balsells, Sara Di Persio, Melanie MY Chan, Matthew Politz, Jann-Frederik Cremers, Nadja Rotte, Sabine Kliesch, Corinna Friedrich, Frank Tuettelmann, Christina Ernst, Sandra Laurentino, Nina Neuhaus, Juan M. Vaquerizas

## Abstract

Gametogenesis, the production of oocytes and sperm, ensures the faithful transmission of genetic material to the next generation. In males, spermatogenesis occurs within the testis, where germ cells progress through a highly ordered developmental programme in close association with supporting somatic cells. Defects in this process cause infertility, yet the gene regulatory mechanisms coordinating normal and dysfunctional human spermatogenesis remain incompletely defined. Here we generate a single-nucleus multiomic and spatial atlas of human spermatogenesis by profiling chromatin accessibility and gene expression in the same nuclei and integrating these data with spatial transcriptomics of intact testicular tissue. We infer high-confidence gene regulatory networks that resolve stage-specific activity of known and novel candidate regulators across germline and somatic compartments, and map spatially restricted signalling interactions within the seminiferous tubule niche. Integration with infertility-associated genetic variation links non-coding risk loci to candidate enhancers and target genes in cell-type-specific regulatory contexts. Finally, profiling clinical cryptozoospermia samples as *in vivo* perturbations supports the ability of this network to capture downstream transcriptional consequences of disease-associated regulatory disruption. Together, these data provide a spatially resolved regulatory framework for human spermatogenesis and a foundation for interpreting the molecular basis of male infertility.

## Introduction

Gametogenesis, the production of haploid oocytes or sperm from diploid precursor cells, is a fundamental process that ensures the faithful transmission of genetic information to the next generation. Defects in this process lead to infertility, which affects an estimated one in six people worldwide, with a male factor implicated in approximately half of these cases ^1,2^. However, only ∼30% of infertile men receive a causal diagnosis ^3^. This diagnostic deficit limits clinical counselling regarding treatment efficacy, the transmission of infertility to offspring, and associated health implications. Furthermore, while assisted reproductive technologies facilitate conception, they are associated with lower pregnancy rates ^4^ and poorer health outcomes in offspring ^5^, indicating an incomplete understanding of the sperm characteristics and associated baseline cellular states in the tissue required for successful reproduction.

To resolve complex disease etiologies in primary and developmental tissues, the integration of single-nucleus RNA sequencing (snRNA-seq), single-nucleus ATAC sequencing (snATAC-seq), and genome-wide association studies (GWAS) provides a robust analytical framework. This multimodal approach has been utilized to map the molecular pathology of Alzheimer’s disease ^6^, coronary artery disease ^7^, and kidney disease ^8–10^ among others. More recently, this framework has expanded to include spatial transcriptomics, allowing dissociated multiomic data to be mapped back onto the native architectural context of complex tissues ^11^. This spatial integration facilitates the resolution of localized cell-cell communication networks ^12^ and the precise functional annotation of enhancer-driven gene regulatory networks (GRNs). Specifically, computational modelling methodologies, such as SCENIC+ ^13^, now enable the direct linkage of chromatin accessibility to transcriptional output, allowing for the mapping of cell-type-specific regulatory control across intact tissue structures.

Within the reproduction field, foundational single-cell transcriptomic and chromatin accessibility studies have mapped cellular heterogeneity during spermatogenesis and gonadal development ^14–32^. However, analysing the male germline presents unique biological challenges, such as incomplete cytokinesis across its differentiation timeline. This leaves developing germ cells connected by intercellular bridges, facilitating extensive transcript sharing ^33–36^. While this feature confounds traditional whole-cell sequencing, single-nucleus isolation intrinsically bypasses this confounder to capture discrete molecular states.

Recently, computational integration of separate single-nucleus ATAC and RNA datasets has been applied to the human testis ^37–39^, but probabilistic label transfer between independent modalities limits the resolution of regulatory networks. Consequently, the cell-type-specific mechanisms of gene regulatory control remain obscure. This limits our understanding of the genetic pathology of male infertility, which is characterized by extreme genetic heterogeneity. While over 500 genes are linked to spermatogenic failure through Mendelian inheritance ^40^, genome-wide association studies (GWAS) have yielded few, poorly replicable risk loci ^41^. Because complex trait-associated variants predominantly reside in noncoding regions, their functional interpretation requires high-resolution maps of cell-type-specific enhancer-promoter interactions. Translating these fragmented genetic associations into mechanistic pathology therefore requires a high-resolution genomic map of transcriptional regulation.

Here, we construct this high-confidence functional map utilizing joint single-nucleus multiomic co-profiling of over 52,000 nuclei from healthy and infertile samples. By capturing chromatin accessibility and gene expression simultaneously, we bypass the computational uncertainty of probabilistic label transfer to generate a directly linked same-nucleus regulatory atlas. From these data, we infer 62 high-confidence eRegulons associated with developmental transitions during human spermatogenesis. We integrate this multiomic network with spatial transcriptomic mapping of over 450,000 cells to orthogonally validate transcriptomic signatures and to resolve spatially-aware cell-cell communication networks. Finally, because the field currently lacks an *in vitro* model system capable of recapitulating complete human spermatogenesis, we utilize clinical infertility samples (cryptozoospermia) as an invaluable *in vivo* perturbation model to assess the predictive power of our healthy reference network.

## Results

### Single nucleus multiomics captures the complexity of the testis

We generated single nucleus multiome (snRNA-seq and snATAC-seq) data from testis biopsies from three men with obstructive azoospermia (OA)—a clinical condition where a physical blockage prevents sperm from reaching the ejaculate, despite completely normal intra-testicular sperm development (Fig. 1A, B) and baseline clinical parameters ^42^ (Supplementary Table S1). After standard quality control and filtering, we retained 19,438 high-quality nuclei for downstream analysis (Fig. S1H), with a mean of ∼5,000 RNA molecules (unique molecular identifiers) and a mean of ∼2,800 ATAC-seq peaks detected per nucleus. RNA and ATAC-seq data were separately processed and then integrated using a weighted nearest-neighbour (WNN) approach ^43^. Based on the expression of known marker genes for different testis cell types ^44^ (Fig. S1B), we identified 14 germline clusters and seven somatic clusters (Fig. 1C, S1A). Four additional clusters could not be assigned to a known cell type. These clusters showed mixed germline-somatic expression (Fig. S1A, B) and/or strongly biased (Fig. S1C) and low (Fig. S1E) RNA/ATAC contributions, and arise from specific replicates (Fig. S1D). These were therefore excluded from further analysis.

**Figure 1:**
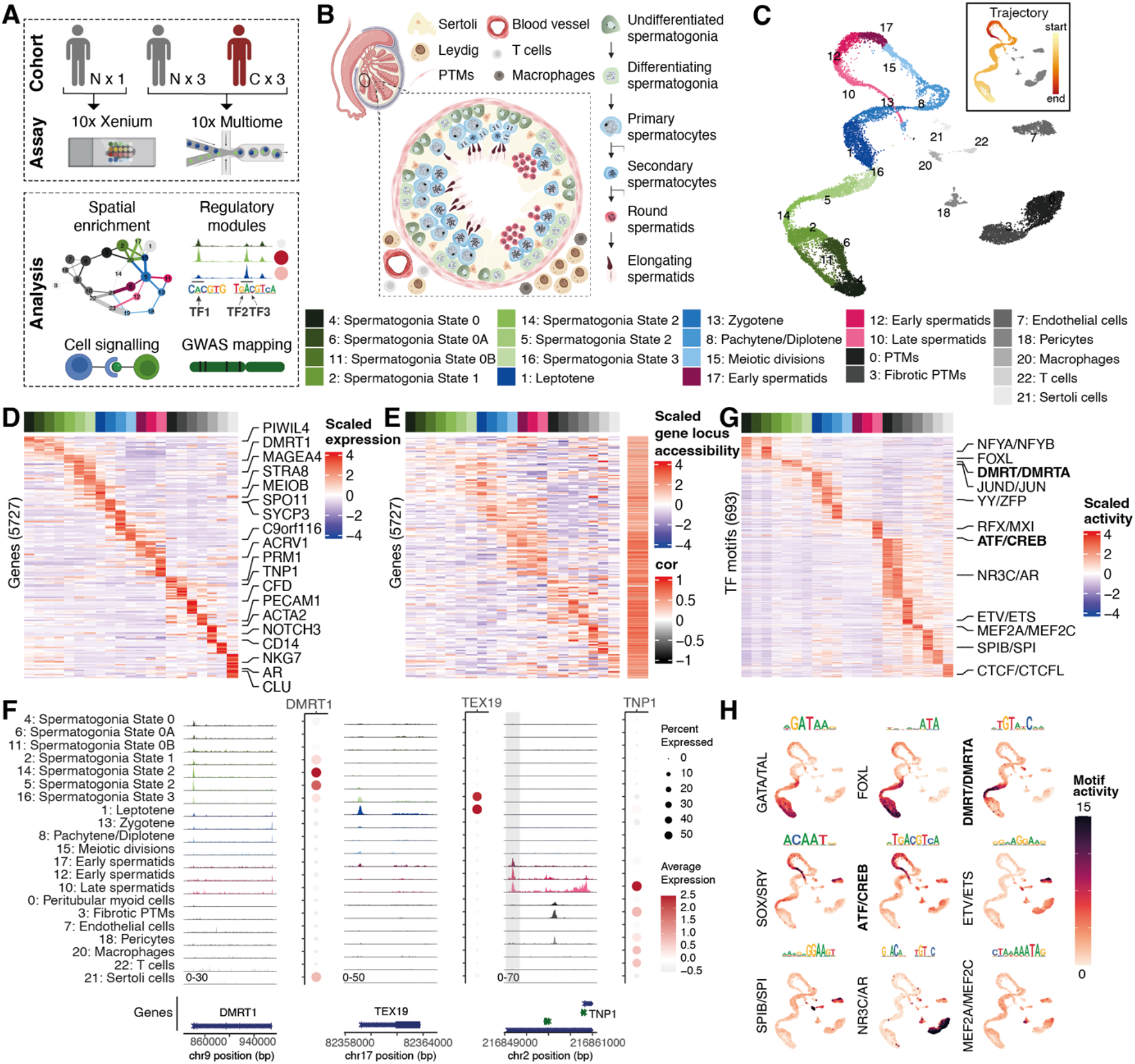
Single-nucleus gene expression and chromatin accessibility characterisation of human spermatogenesis. A. Schematic overview of the study design. 3 samples with healthy spermatogenesis, 3 with cryptozoospermia were processed with 10x multiome technology and 1 sample with predominantly tubules with full spermatogenesis was processed with Xenium spatial technology. Healthy samples were obtained from patients diagnosed with obstructive azoospermia (OA), a condition characterized by the absence of spermatozoa in the ejaculate despite preserved spermatogenesis ^42^. B. Schematic overview of the stages of human spermatogenesis. Undifferentiated spermatogonia commit to differentiation, then become spermatocytes which go through meiosis to produce haploid spermatids. The seminiferous tubules contain germline and Sertoli cells, are surrounded by peritubular myoid cells (PTMs), and the interstitial space between tubules contains Leydig cells, pericytes, endothelial cells, and immune cells. Seminiferous tubules of different stages include different subsets of germline developmental stages; the schematic tubule shown here includes segments of different stages in order to show all germline cell types. C. UMAP representation of single-nucleus multiomics data from human testis. Inset shows trajectory inference performed on germline clusters using Slingshot. D. Heatmap of gene expression of top 500 protein-coding genes per cluster (in total 5727 genes) across spermatogenesis. Germline clusters are ordered by their position in the trajectory shown in B, followed by somatic cells. Genes are ordered by the cluster in which the gene has max expression. Selected marker genes for different cell types are labelled on the right-hand side. E. Heatmap of the accessibility at gene locus (comprising the gene body and the 2 kb upstream putative promoter region) of top 500 protein-coding genes per cluster shown in C across spermatogenesis. Germline clusters are ordered by their position in the trajectory shown in B, followed by somatic cells. Accessible regions are ordered by the cluster in which the peak has max accessibility. Pearson correlation between expression and locus accessibility across the clusters for the top 500 protein-coding genes shown in C. F. Examples of dynamic chromatin accessibility at selected marker genes. Each sub-panel shows cluster-specific pseudobulk chromatin accessibility tracks on the left, with gene annotations below. Regions are centred on genes of interest, which are labelled. Expression of the marker gene of interest in the cluster is shown in the dot plots on the right, where colour indicates scaled average expression and size indicate the % of cells in the cluster with detected expression. The *TNP1* locus contains a highly accessible putative enhancer (highlighted in grey) in addition to gene-associated accessibility. G. Heatmap of motif activity of 693 TF motifs across spermatogenesis. Germline clusters are ordered by their position in the trajectory shown in B, followed by somatic cells. Motifs are ordered by the cluster in which the motif has highest activity. The transcription factor families for selected cluster-specific motifs are shown on the right-hand side. H. UMAP embedding of motif activity across single cells for selected TF motifs highlighting stage-specific motif activity.

Importantly, the chromatin accessibility data allowed us to further resolve cluster identities that were unclear based on expression data alone. Cluster 15 has low expression of most marker genes (Fig. S1B). However, we observed that this cluster exhibits a bimodal distribution of ATAC-seq reads from chromosome X, indicating the separation of cells into chromosome X- or Y-bearing cells after meiosis I (Fig. S1O), along with the expression of *C9orf116*, a previously identified marker of secondary spermatocytes ^15^. We therefore considered cluster 15 to represent the rare population of cells undergoing meiosis II and not yet expressing spermatid markers. Among the somatic cells we could identify distinct clusters representing Sertoli cells, PTMs, endothelial cells, pericytes, and testicular immune cells. In comparison with previous scRNA-seq datasets, we identified a lower proportion of spermatids and we do not identify a distinct cluster of Leydig cells until integrated and clustered with the published datasets ^16,26^ (Fig. S2A, B). In contrast, we identify a larger proportion of spermatogonia (Fig. S2A, B) likely reflecting variation in nuclei isolation efficiency across cell types. Importantly, corresponding cell types show high transcriptional concordance with published datasets (Fig. S2C), supporting the robustness of our dataset and cell-type annotations.

Trajectory inference using Slingshot ^45^ recapitulates the known continuum of spermatogenesis with a main trajectory that includes more than 99% of germline cells (inset, Fig.1C). Clusters 13: Zygotene and 8: Pachytene/Diplotene show the highest number of expressed genes and accessible peaks and average counts per gene and peak (Fig. S1I, J, L, M). This increased chromatin opening may be linked to widespread expression in these stages ^29,46^ and may also reflect changes in chromatin state that precede replacement of histones by protamines ^47^. Differential gene expression analysis identified a total of ∼12,000 genes displaying significant positive enrichment across the individual clusters, including many known markers for the different cell types in the testis (Fig. 1D, F, Supplementary Table S2). Examining chromatin accessibility at these loci shows dynamic promoter and gene body accessibility across the different cell types which correlates with gene expression (Fig. 1D, E and Fig.S1P, S1G, P). Approximately 65% of these genes (3,721/5,727) show a significant Pearson correlation between expression and locus accessibility, of which 99.5% (3,703/3,721) are positively correlated (median *r* = 0.7). In addition, we also identify distal peaks with stage-specific accessibility during spermatogenesis, which represent putative enhancers. An example is shown at the *TNP1* locus, where a peak upstream of the gene gains accessibility in early spermatids (Fig. 1F). In all cell types, promoters generally exhibited higher baseline accessibility compared to dynamic distal peaks (Fig. S1L, M).

However, promoter accessibility showed a lower correlation with steady-state gene expression compared to gene bodies (Fig. S1P). Our data indicate that promoter accessibility and transcriptional activity are not consistently coupled throughout spermatogenesis. We observe that chromatin often maintains a broadly permissive state across multiple lineages, even when transcriptional output is highly restricted. Conceptually mirroring recent developmental findings ^48^, this extensive epigenetic priming may help explain why transcriptome-only analyses incompletely resolve cis-regulatory relationships during spermatogenesis.

To understand how these changes in chromatin accessibility might be regulated, we examined changes in accessibility for the motifs of 693 transcription factors genome-wide ^49^, revealing clear cluster-specific patterns of motif activity that identify putative regulators of key transitions during spermatogenesis (Fig. 1G, H, Supplementary Table S3). Importantly, these *in silico* predictions closely mirror known *in vivo* biological functions. For example, we observe high motif activity for DMRT1 specifically in spermatogonia, consistent with its essential role in early germ cell maintenance ^50^. In contrast, motif activity for CREM is strikingly induced in post-meiotic spermatids, perfectly aligning with its known function as a master transcriptional driver of spermiogenesis ^51,52^. By capturing these dynamic regulatory transitions, our multiome data provides a robust framework for mapping the stage-specific control of human spermatogenesis.

Beyond localized regulation, our chromatin accessibility data also capture the silencing of unsynapsed regions of the sex chromosomes during meiosis (meiotic sex chromosome inactivation; MSCI), shown by the global decrease in signal on the X chromosome in pachytene/diplotene compared to autosomes (Fig. S1N, O). In addition, we observe a marked drop in the fraction of ATAC-seq reads falling in discrete peaks after meiosis consistent with global post-meiotic silencing of the genome (Fig. S1K). Altogether, our single nucleus multiome atlas captures the gene expression and regulatory dynamics of a comprehensive repertoire of germ cells throughout spermatogenesis, and of their supporting somatic cells.

### Spatial transcriptome captures complexity of the testis

While single-nucleus multiomics resolves discrete regulatory states, tissue dissociation inherently obscures their native spatial context. To map these cellular profiles back to the intact microenvironment of the seminiferous tubule, we generated spatial transcriptomic data on a morphologically normal testicular tissue section obtained adjacent to a germ cell tumour (60.6 mm²), utilizing the 10x Xenium In Situ platform with a custom 500-gene panel derived from our sn-multiome data. Cell segmentation was performed using the Xenium Onboard Analysis v2.0 pipeline with a 5 µm nuclear expansion to define cellular boundaries. After quality control, 451,388 cells were retained, comprising a mean of ∼129 RNA molecules (UMIs) detected per cell, across 1,325 seminiferous tubule cross-sections.

Using marker gene expression together with RCTD-based label transfer (S3A, B), we identified 11 germline clusters and 12 somatic clusters. Within the tissue, two types of seminiferous tubules were observed, which we term full (n = 1,116) and empty (n = 209), based on hierarchical clustering of tubules according to cell-type composition (Fig. S3G, H). Full tubules contain the complete spectrum of germ cells across developmental stages together with supporting somatic cells (Fig. 2A-E, S3B). In a representative full tubule (Fig. 2A-E), peritubular myoid cells (PTMs) form the outer wall of the tubule. Within the tubule, Sertoli cells and germ cells—including spermatogonia, meiotic cells at different stages, and spermatids—are arranged from the periphery toward the lumen, recapitulating the architecture of a healthy adult testis. In contrast, empty tubules retain spermatogonia and supporting somatic populations but lack post-spermatogonial germ-cell stages (Fig. S3C-G). These tubules localized adjacent to the tumour margin and expressed the Germ Cell Neoplasia in Situ (GCNIS) marker OCT4 ^53^ (Fig. S3E), indicating a localized pathological transformation of normal spermatogenesis. Consequently, only full tubules were utilized for downstream analyses of healthy spermatogenesis.

**Figure 2:**
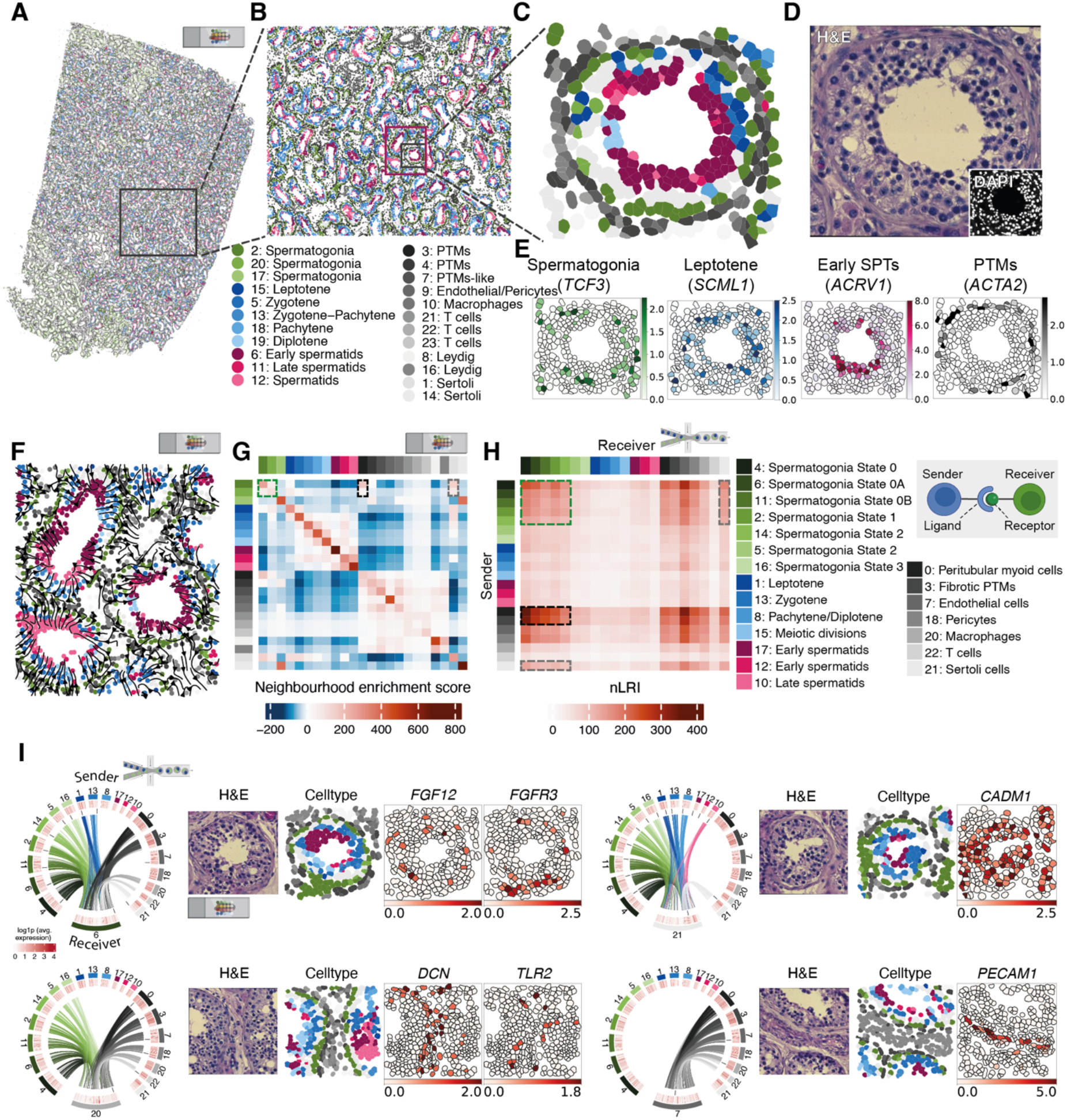
Single cell spatial transcriptomic characterisation of human spermatogenesis. A. Scatterplot of single-cell spatial coordinates obtained from 10x Xenium In Situ platform of a human testicular tissue cross-section of 60.6 mm². Each point represents an individual cell positioned according to its spatial location within the tissue section and coloured based on celltype cluster assignment. Slide and microfluidics icon indicate origin of data as spatial and multiome respectively. B. Zoom-in on a section of A focussing on a collection of full tubules. C. Zoom-in on a section of B focussing on a single full tubule. D. H&E staining (and inset, DAPI staining) of the full tubule in C. E. Marker gene expression for *TCF3* representing spermatogonia, *SCML1* for leptotene, *ACRV1* for early spermatids and *ACTA2* for PTMs in the full tubule in C. F. Predicted cellular trajectories inferred by SpaTrack, within the pink box highlighted in B, based on gene expression and spatial information from the xenium data and projected onto the tissue section with cell areas coloured based on cell type cluster assignment. Flow arrows indicate the inferred direction of cell-state transitions across spatial locations. G. Heatmap of neighbourhood enrichment score inferred from the spatial location of the different cell types in the spatial data. Rows and columns are arranged in developmental order with the germ cells followed by the somatic cells. Dashed boxes highlight positive enrichment score between spermatogonia-spermatogonia (green), spermatogonia-PTMs (black) and spermatogonia-Sertoli cells (grey). H. Heatmap of number of LRIs between sender and receiver cell types inferred from the multiome data. Rows and columns are arranged in developmental order with the germ cells followed by the somatic cells. Dashed boxes indicate high number of interactions between the cell type pairs highlighted in C) including spermatogonia-spermatogonia (green), spermatogonia-PTMs (black) and spermatogonia-Sertoli cells (grey). Inset represents a schematic of ligand-receptor interaction between sender and receiver cells. I) Circos plots for selected clusters representing significant LRIs (BH adjusted p.value < 0.01) between sender and receiver cell types inferred from the multiome data for cell type pairs with positive neighbourhood enrichment score in the spatial data. The outer-most track represents the cell type, followed by a track of heatmap of gene expression for the ligand on the sender cell types and for the receptor on the receiver cell types in the multiome data. The third track indicates LRIs for which the ligand and receptor gene expression is represented in the spatial data on the right. The inner-most track represents links between the ligand receptor pairs, with the thickness of edge indicating strength of interaction and the colour indicating the sender cell type.

Within full tubules, trajectory analysis integrating gene expression and spatial coordinates using SpaTrack ^54^ reconstructed the developmental progression from spermatogonia to spermatids (Fig. 2F, pink box in Fig. 2B). Representative spatial trajectories map progression either up to the meiotic stage or through to haploid spermatids (Fig. S3I). Cellular neighbourhood quantification revealed significant spatial clustering of spermatogonia with themselves (green box), peritubular myoid cells (PTMs, black box), and Sertoli cells (grey box) (Fig. 2G), quantitatively confirming their established architectural niche.

To determine how these physical neighbourhoods dictate intercellular signalling, we analysed ligand–receptor interactions (LRIs) using scDiffCom ^55^. We first identified 36,224 significant LRIs in the multiome dataset (Supplementary Table S4). Consistent with spatial proximity, the highest interaction frequencies occurred between physically neighbouring cell types, particularly spermatogonia with PTMs and Sertoli cells (Fig. 2H). To eliminate biologically improbable interactions, we filtered this multiomic cell-cell communication network using the spatial neighbourhood data, retaining only LRIs between cell-type pairs with positive spatial neighbourhood scores. This spatially constrained network yielded 21,367 interactions with distinct localized signalling patterns (Fig. 2I, S4A, B).

Spatially-informed interactions resolved distinct signalling compartments within and outside the tubule. Within the basal niche, State 0A spermatogonia (cluster 6) act as a primary signalling hub, receiving signals from multiple spermatogonial clusters, leptotene–zygotene cells, PTMs, immune cells, and Sertoli cells (Fig. 2I, top-left panel). Key spatial maps highlighted localized FGF12–FGFR3 signalling, where FGF12 expressed by neighbouring somatic and early meiotic cells targets FGFR3 on State 0A spermatogonia (Fig. 2I, top-left panel). Similarly, we identified NLGN1–NRXN1 signalling specifically between the spermatogonial compartment and PTMs, as well as DCN–ERBB4 signalling linking PTMs to developing germ cells (Fig. S4B).

Beyond the spermatogonial niche, spatial maps highlighted structural and immune signalling networks. CADM1–CADM1 homotypic interactions were abundant between Sertoli cells (cluster 21) and germ cell populations, with high CADM1 expression definitively localized to Sertoli cells, as also recently reported in ^23^, alongside germ cells (Fig. 2I, top-right panel).

Finally, extratubular somatic populations, such as endothelial cells (cluster 7), primarily received signals from outside the seminiferous tubule. This included extensive DCN–TLR2 signalling between PTMs, pericytes, endothelial cells, and macrophages (cluster 20) (Fig. 2I, bottom-left panel), as well as PECAM1 homotypic interactions mapping exclusively to endothelial and macrophage populations (Fig. 2I, bottom-right panel). Together, this spatially restricted interactome defines the localized signalling networks maintaining human spermatogenesis.

### Construction of a testis gene regulatory network

To characterize the gene regulatory landscape underlying human spermatogenesis, we utilized SCENIC+ to integrate chromatin accessibility, gene expression, and TF motif enrichment ^13^. This approach predicts candidate cis-regulatory elements, upstream regulatory transcription factors (TFs), and target genes, collectively termed an eRegulon (Fig. 3A). We identified 62 high-confidence eRegulons exhibiting strong correlations between TF expression, predicted enhancer accessibility, and target gene expression. These comprise 50 activating eRegulons (targeting an average of 1,250 regions and 620 genes) and 12 repressive eRegulons (targeting an average of 300 regions and 150 genes).

**Figure 3:**
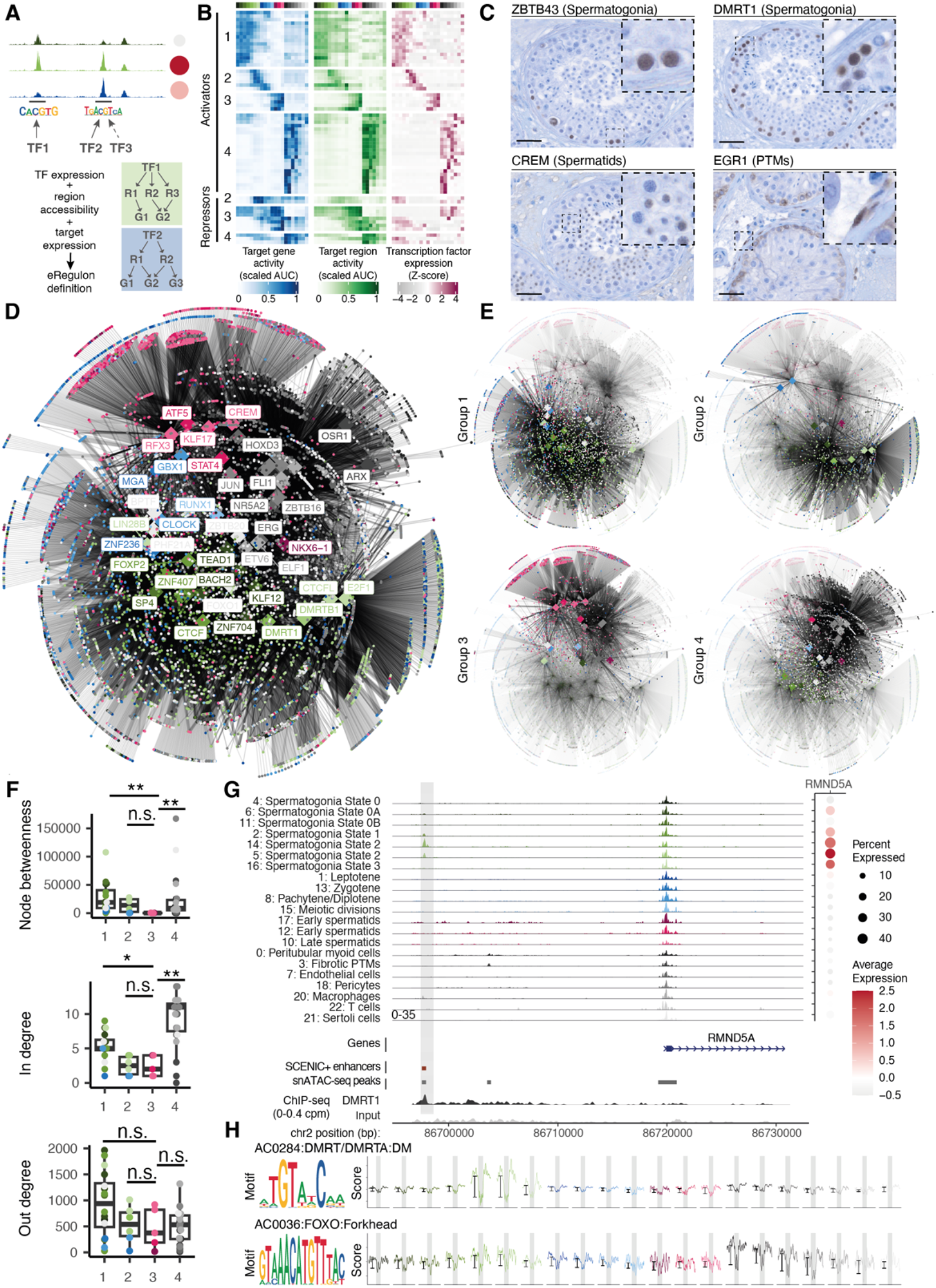
SCENIC+ identifies stage-specific eRegulons and indicates differences in regulation between spermatids and other stages. A. Schematic depiction of eRegulon identification by SCENIC+. SCENIC+ identifies putative enhancers containing transcription factor (TF) motifs. TFs, putative enhancer regions (R1-R3), and target genes (G1-G3) are grouped into eRegulons on the basis of correlation between the TF expression, target gene expression, and region accessibility. An eRegulon consists of a TF, its target regions, and its target genes. B. 62 high-confidence eRegulons which show dynamic activity across cell types were identified. Top, activator eRegulons. Bottom, repressor eRegulons. Left-to-right, panels represent eRegulon target gene expression enrichment per cluster (blue, scaled average AUC), eRegulon target region accessibility enrichment per cluster (green, scaled average AUC), and TF expression per cluster (magenta, Z-score). There are 4 distinct groups of eRegulons active in spermatogonia (1), differentiating spermatogonia and spermatocytes (2), spermatids (3) and somatic cell types (4). C. Representative immunohistochemical staining of human seminiferous tubule sections showing expression of the candidate eRegulon transcription factors ZBTB43 and DMRT1 in spermatogonia, CREM in spermatids, and EGR1 in peritubular myoid cells (PTMs). Nuclei were counterstained with hematoxylin. Scale bars, 40μm. D. The gene regulatory network made up of the 62 identified high-confidence eRegulons. Nodes represent genes, coloured by the cluster in which the gene has max expression. Colour assignments are as in Fig.1. TFs are labelled and shown as large diamonds and all other genes as smaller circles. (Note: some TF text labels are graphically suppressed to prevent overlap, but all 56 TFs are plotted as nodes). Label background colour is white or grey for readability purposes only. Edges represent a regulatory relationship between a TF and a target gene inferred by SCENIC+. E. Representative subnetworks from the gene regulatory network shown in D, highlighting regions corresponding to the eRegulon groups defined in B. Node and edge annotations are as described in D. F. Quantification of network properties for activating TFs. TFs driving eRegulons in Group 3, active in spermatids, have significantly lower betweenness centrality and in degree compared to TFs active in Group 1 or Group 4. Significance asterisks are shown for pairwise comparisons of Group 3 with all other groups; **: p <= 0.01, *: p <= 0.05, ns: p > 0.05. G. SCENIC+ identifies an enhancer (red bar within grey highlight) upstream of *RMND5A* which is regulated by DMRT1, DMRTB1, and FOXO1. Left, cluster-specific pseudobulk chromatin accessibility tracks; right, *RMND5A* expression shown as a dotplot where colour indicates scaled average expression and size indicates the % of cells in the cluster with detected expression. Bottom, gene annotations, predicted enhancers (red), and scATAC-seq peaks (grey). A DMRT1 ChIP-seq peak overlapping the predicted enhancer is shown with a blue bar. H. TF footprinting for motifs bound by DMRT1/DMRTB1 (top) and FOXO1 (bottom), showing strong binding in differentiating spermatogonia.

Because several TFs act as both activators and repressors depending on regulatory context, these eRegulons comprise a total of 56 unique TFs that are linked to over 10,000 unique genes and 23,000 accessible regions, including ∼12,000 distal predicted enhancers. Based on target gene activity, these eRegulons form four clusters: those active primarily in undifferentiated spermatogonia (Group 1; 16 eRegulons), differentiating spermatogonia and/or spermatocytes (Group 2; 12 eRegulons), spermatids (Group 3; 8 eRegulons), and somatic cells (Group 4; 28 eRegulons) (Fig. 3B, S6A). Known stage-specific regulators were recovered in the expected groups, including DMRT1 (Group 2) and CREM (Group 3) (Fig. S6A).

We further validated candidate TF expression at the protein level via immunohistochemistry, confirming ZBTB43 and DMRT1 in spermatogonia, CREM in spermatids, and EGR1 in PTMs (Fig. 3C) using histologically normal testis sections (Supplementary Table S5). To benchmark our distal regulatory predictions against external functional data, we cross-referenced our SCENIC+-predicted elements with recently reported in vitro human testicular enhancer assays ^25^. This comparison revealed modest but biologically informative overlap, including concordant signal at several loci with known roles in spermatogenesis, such as PRDM9 (Fig. S5). At the same time, we noted that a substantial fraction of previously tested regions corresponded to promoter-proximal accessible elements rather than distal enhancers (Fig. S5C-E) or did not overlap accessible regions in our *in vivo* atlas (Fig. S5H-N). These discrepancies likely reflect both methodological differences between *in vivo* multiomic mapping and *in vitro* reporter assays, and the difficulty of establishing which sequences function as bona fide distal regulatory elements outside the native context of adult human testis.

Integrating these 62 eRegulons generated a global gene regulatory network of the human testis (Fig. 3D, E). Network topology revealed that TFs active in somatic cells and spermatogonia localized to the dense central core, whereas spermatid-active TFs occupied the periphery. We quantified this using betweenness centrality—a measure of a TF’s frequency on pathways linking other gene pairs. TFs active in spermatids (Group 3) exhibited significantly lower betweenness centrality than those in undifferentiated spermatogonia (Group 1) and somatic cells (Group 4) (Fig. 3F). Degree analysis indicated this architecture is at least partially driven by spermatid-specific TFs possessing lower in-degrees, whereas spermatogonial TFs exhibit high in- and out-degrees, signifying extensive cross-regulation during early differentiation.

Within the germline compartment, SCENIC+ successfully resolved stage-specific regulatory activation. For example, the E3 ubiquitin-protein transferase RMND5A exhibits maximal expression in differentiating spermatogonia (State 2). We identified a distal candidate enhancer for RMND5A predicted to be regulated by DMRT1, DMRTB1, and FOXO1.

Providing strong orthogonal validation for our multiomic network, this predicted enhancer localizes to experimentally confirmed DMRT1 binding sites in healthy human testis ^56^ (Fig. 3G). TF footprinting using TOBIAS ^57^ confirmed DMRT1 and FOXO1 activity specifically in differentiating spermatogonia, coincident with both enhancer accessibility and *RMND5A* expression (Fig. 3H). Genome-wide, DMRT1 ChIP-seq peaks showed a highly significant overlap with SCENIC+-predicted *DMRT1* enhancers (27.6% vs. 2.6% background, permutation test p = 0.001). Similarly, we resolved distinct temporal regulation for *RNF17*, identifying two candidate enhancers: one active in undifferentiated spermatogonia (driven by LIN28B) and a second active in zygotene/pachytene (activated by LIN28B and CLOCK; repressed by JUN), supported by robust TF footprinting (Fig. S6B, H). Notably, this distinct bi-phasic expression pattern in human spermatogenesis diverges from the expression profile of the mouse ortholog (Rnf17), which is predominantly restricted to post-meiotic spermatids ^58^. This highlights the critical necessity of human-specific regulatory mapping to accurately model clinical pathology. Furthermore, we mapped celltype-specific regulation for *VGLL3*, identifying one enhancer driving spermatid expression via ATF5/CREM, and a distinct enhancer driving PTM expression via EBF1/NR2F2 (Fig. S6C, H).

Within the somatic compartment, this multiomic network captured the regulatory programs maintaining the structural and immunological niche. In PTMs, we identified seven candidate enhancers for *DCN* predicted to be activated by FOS, JUN, MAF, MYC, NR2F2, OSR2, and ARX, and repressed by ERG and RUNX1, supported by strong TF footprinting (Fig. S3D, H). In Sertoli cells, SCENIC+ mapped enhancers activating *DEFB119* via ELF1 binding, a regulatory link supported by robust localized footprints (Fig. S3E, H). Finally, within the extratubular somatic populations, we identified specific enhancer networks regulating *ARHGAP15* (via ELF1) in immune cells (Fig. S3F, H) and the efflux transporter *ABCB1* (via ETS2, FLI1, and KLF2) in endothelial cells (Fig. S3G, H). The full compendium of high-confidence TF-target gene pairs is provided in Supplementary Table S6.

Collectively, these data delineate the cell-type-specific regulatory architecture governing healthy human spermatogenesis, providing a foundational reference for evaluating the functional impact of noncoding genetic variants associated with male infertility.

### The spermatogenesis gene regulatory network identifies targets of infertility-associated variants

Genome-wide association studies (GWAS) for complex traits frequently identify non-coding variants, which are enriched within cell-type-specific candidate cis-regulatory elements. To date, five GWAS of male infertility have identified 14 independent Single Nucleotide Variants (SNVs) associated with disease in Han Chinese or European populations (Supplementary Table S7) ^41,59–62^. Because most (13) of these SNVs reside in non-coding regions, their mechanistic interpretation remains unresolved. We therefore leveraged our multiomic network to map these variants to their potential target genes.

Given none of the 13 non-coding lead SNVs directly overlapped a SCENIC+-predicted enhancer in our dataset, we expanded our analysis to include SNVs in high linkage disequilibrium (LD) with the lead variants (LD; r^2^ > 0.8) using 1000 Genomes Project Phase 3 data ^63,64^ (Fig. 4A, inset). To capture candidate linked variants that might include causal alleles co-inherited with the lead variants ^65^, we assessed regions within 500 kb of each lead variant, identifying between 2 and 124 additional SNVs. Notably, rs16824398 (in LD with lead variant rs2477686, r^2^ = 1) directly overlapped a predicted enhancer within the *PLCH2* gene body. This candidate enhancer is highly accessible in PTMs and pericytes, and is predicted to be regulated by FOXC1, EBF1, and NFIC to target two genes: *MMEL1* (an endopeptidase essential for normal sperm function ^66,67^) and *TNFRSF14* (a member of the TNF receptor superfamily previously associated with azoospermia ^68^) (Fig. 4A-D).

**Figure 4:**
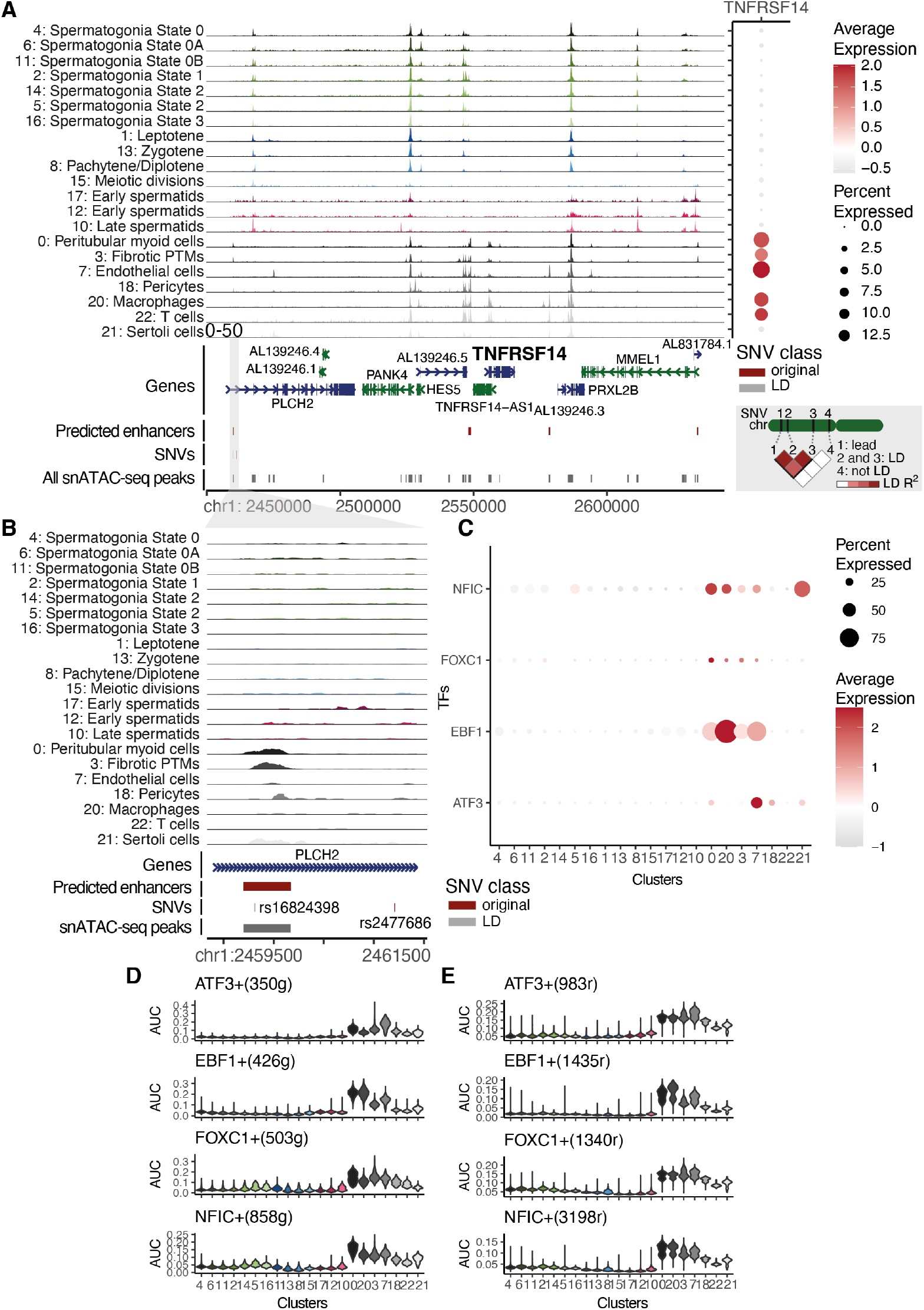
SCENIC+ results identify TNFRSF14 as a candidate target gene for a GWAS locus associated with infertility. A. *TNFRSF14* locus. Left, cluster-specific pseudobulk chromatin accessibility tracks. Right, *TNFRSF14* expression per cluster as a dotplot; colour indicates scaled average expression and size indicates the % of cells in the cluster with detected expression. Bottom, gene annotations, SCENIC+ predicted distal enhancers for *TNFRSF14*, SNVs, and snATAC-seq peaks. The lead SNV (dark red) is rs2477686 identified as associated with male infertility by Hu et al. 2012 ^59^; variants in high LD (r^2^ > 0.8) with this SNV are shown in grey. Inset shows a schematic of linkage disequilibrium (LD). B. Top, close up of part of the region in A. rs16824398, a variant in high LD with rs2477686 overlaps an snATAC-seq peak identified by SCENIC+ as a putative enhancer for *TNFRSF14*, regulated by ATF3, EBF1, FOXC1, and NFIC. C. Expression of *ATF3*, *EBF1*, *FOXC1*, and *NFIC* as a dotplot; colour indicates scaled average expression and size indicates the % of cells in the cluster with detected expression. D, E. eRegulon activity for ATF3, EBF1, FOXC1, and NFIC across clusters, as measured by target gene expression enrichment (D) and target region chromatin accessibility enrichment (E) by their corresponding AUC (Area Under Curve) scores.

Furthermore, two additional variants in high LD with rs2477686 (rs2494631 and rs2494632, R^2^ = 1) overlapped a promoter-proximal regulatory region active in spermatids. While excluded from the final distal-enhancer SCENIC+ set due to its promoter proximity, this region also regulates *MMEL1*. This mapping is consistent with the enriched expression and eRegulon activity of its predicted upstream regulator, KLF17, in spermatids (Fig. S7C-E).

Strict genome-wide significance thresholds in highly heterogeneous traits like male infertility often obscure functionally relevant polygenic signals. To identify sub-threshold variants that might drive disease risk through specific gene regulatory networks, we intersected our multiomic data with suggestive SNVs (p < 10^-5^) from a recent male infertility meta-analysis of 10,886 cases and 995,982 controls ^41^, resulting in a total of 504 suggestive SNVs and 897 SNVs in high LD. Three suggestive SNVs overlapped SCENIC+-predicted enhancers, allowing us to map their cell-type-specific regulatory wiring (Fig. S7A). First, rs114935822 mapped to a PTM- and pericyte-accessible enhancer regulating *SORBS1*—an adaptor protein upregulated in teratospermia models ^69^—via AR and ZEB1 (Fig. S7F-H). Second, rs749792650 overlapped an enhancer regulating *APOL1* and *APOL2*, lipid-binding proteins linked to innate immunity ^70–72^ and non-obstructive azoospermia ^73^ respectively. This locus is broadly accessible across PTMs, pericytes, and macrophages, but its predicted transcription factors (including BCL6B, ERG, and ETS1) show maximal expression and eRegulon activity in endothelial cells (Fig. S7I-K). Finally, rs74863147 mapped to an enhancer predicted to regulate *SYNJ2*. While SYNJ2 has an established role in spermatid manchette formation ^74,75^, the chromatin accessibility of this specific regulatory element is restricted to endothelial cells, where *SYNJ2* is also expressed (Fig. S7L-N).

While mapping individual GWAS loci to specific SCENIC+ enhancers yields targeted mechanistic hypotheses, male infertility is a highly polygenic trait. To evaluate the global contribution of the broader cis-regulatory landscape to disease heritability, we expanded our scope from the highly stringently defined eRegulons to the complete genome-wide set of cell-type-specific ATAC-seq peaks. We utilized stratified linkage disequilibrium score regression (S-LDSC ^76^) to test whether variants within these adult human testis open chromatin regions contribute disproportionately to the heritability of male infertility. Using publicly available European male infertility GWAS summary statistics (6,885 cases and 410,578 controls from ^41^), we observed positive coefficient estimates across several stages of spermatogenesis—most notably in zygotene, pachytene, and diplotene spermatocytes— although none of these enrichments remained statistically significant following FDR correction (Fig. S7B).

Collectively, mapping these loci against our high-resolution network transforms both genome-wide significant and suggestive SNV associations into localized, testable regulatory hypotheses. Given the current statistical power limitations of male infertility GWAS, this integrated framework provides a reference for prioritizing putative non-coding variants and evaluating their functional impact on the spermatogenic regulatory landscape.

### Gene regulatory network perturbation in infertility

Given the limitations of *in vitro* spermatogenesis models ^77^, we utilized clinical samples to test whether the inferred network captures disease-associated regulatory perturbation *in vivo*. We performed single-nucleus multiomics on testicular biopsies from three individuals with cryptozoospermia (sperm concentration < 0.1 million/ml^3^) (Supplementary Table S8). This yielded 32,583 high-quality cells, with a mean of ∼3,300 RNA molecules (UMIs) and ∼2,200 ATAC-seq peaks detected per cell. By computationally integrating this patient data with our normal controls, we anchored the diseased cells to a healthy reference manifold, enabling precise, comparative cell-state mapping. Following integration, we resolved all expected germline and somatic lineages, including Leydig cells (Fig. 5A, S8A). Visually separating the cohorts allows for immediate appreciation of the phenotypic differences: while all three patients shared a severe depletion of post-meiotic spermatids (Fig. 5A, B), their underlying molecular profiles were highly heterogeneous, exhibiting minimal overlap in differentially expressed genes (DEGs) or differentially accessible (DA) peaks (Fig. 5C, S8B-E). Highlighting this distinct aetiology, exome sequencing identified a pathogenic variant in M1AP—a known regulator of meiotic recombination—exclusively in Crypto Rep 1 ^78,79^.

**Figure 5:**
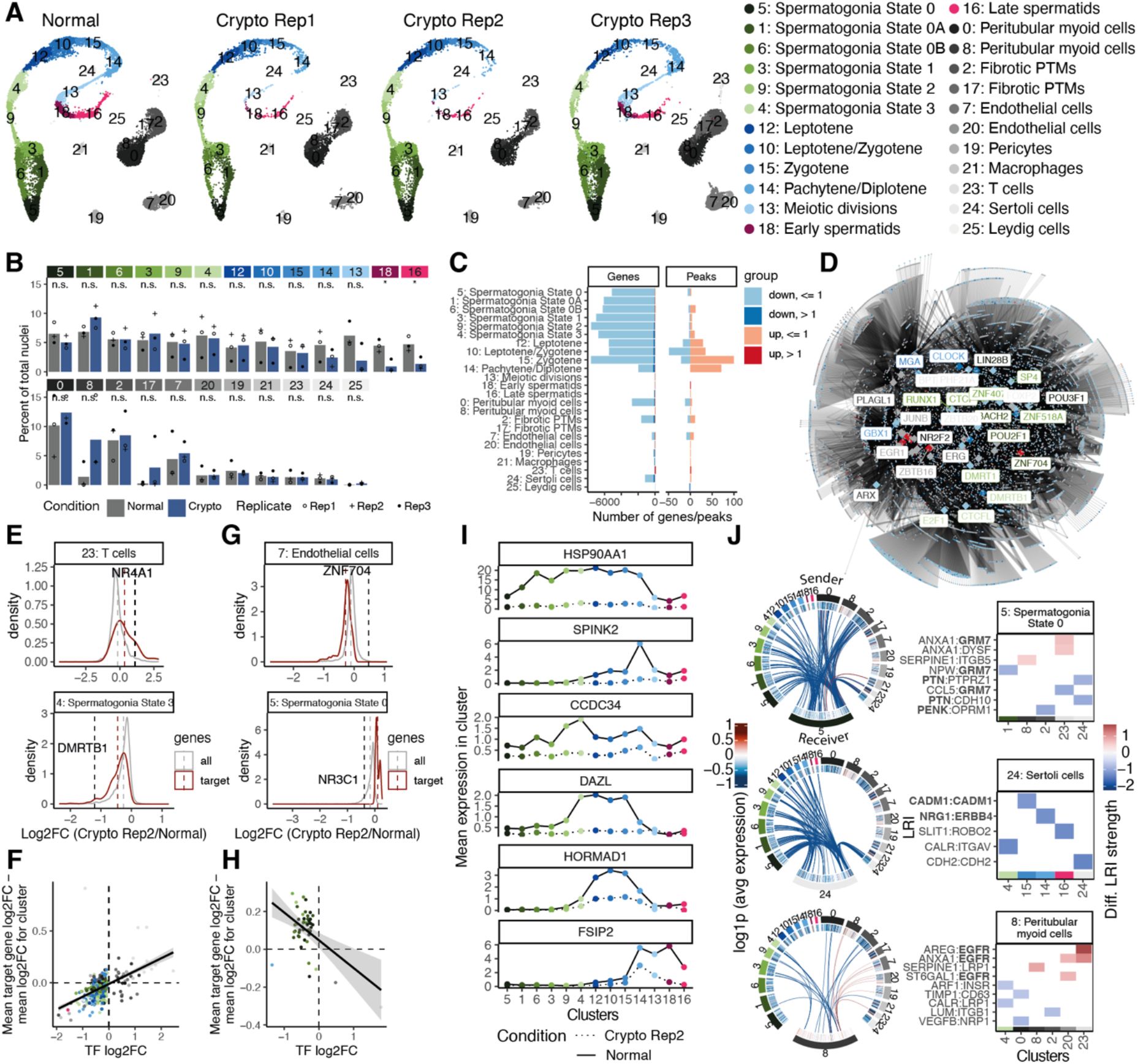
Single-cell multiomics of spermatogenesis in infertile men. A. UMAP representation of single-nucleus multiomics data from normal spermatogenesis (left, combined samples), and two infertile individuals with cryptozoospermia. B. Quantification of the contribution of each biological replicate to the clusters shown in A. The samples from infertile individuals have a significant depletion in the number of spermatids (clusters 18 and 16), and a noticeable increase in one subcluster of PTMs (cluster 8) and fibrotic PTMs (cluster 17). C. Numbers of differentially expressed genes (left) and differentially accessible peaks (right) in each cluster, comparing normal spermatogenesis and Crypto 2. D. Differentially expressed genes between normal spermatogenesis and Crypto 2 shown in the context of the high-confidence gene regulatory network (Fig. 3D). TFs are shown as large diamonds and all other genes as smaller circles. Genes that are significantly downregulated or upregulated in any cluster are shown in blue or red respectively and all other genes in grey. Edges represent a regulatory relationship between a TF and a target gene inferred by SCENIC+. Differentially expressed TFs are labelled and coloured according to the cluster in which they have highest expression in the normal spermatogenesis samples. E. The target genes of significantly differentially expressed SCENIC+ TFs show changes in gene expression consistent with SCENIC+ predictions. For positive eRegulons, examples of the distribution of log2FC in expression between Crypto 2 and normal samples, for all genes (grey) and NR4A1 target genes (red, top) and DMRTB1 target genes (red, bottom) representing an upregulated and a downregulated activator TF respectively. The mean of the distribution is shown with a corresponding vertical dashed line. The log2FC of NR4A1 or DMRTB1 is shown with a black vertical dashed line. The difference in mean log2FC between target genes and all genes is plotted on the y axis in panel F. F. Activator TFs significantly differentially expressed in any cluster between normal and Crypto 2 (p < 0.05) are shown. Each point represents a TF and its targets in a given cluster; the logFC of the TF is represented on the x axis and the difference in mean log2FC between target genes and all genes is plotted on the y axis, as explained in panel E. Clusters are coloured according to the key in A. Points with a Fisher test p-value of < 0.05 for the association between direction of change in expression for the TF and direction of change in expression of its targets are shown G. Same as E, for negative eRegulons, ZNF704 (top) and NR3C1 (bottom) are shown as examples of an upregulated and downregulated repressor TF respectively. The difference in mean log2FC between target genes and all genes is plotted on the y axis in panel H. H. Same as F, for repressor TFs. I. Examples of genes that are downregulated in Crypto 2 compared to normal spermatogenesis samples. Average normalised expression per cluster is shown for normal (solid line) and Crypto 2 (dotted line). Colours correspond to germline clusters according to the key in panel A. J. Circos plots representing significantly differential LRIs (BH adjusted p.value < 0.01) between sender and receiver cell types inferred from the multiome data for cell type pairs with positive neighbourhood enrichment score in the spatial data. The outer-most track represents the cell type, followed by a track of heatmap of log2FC of gene expression between crypto and healthy conditions for the ligand on the sender cell types and for the receptor on the receiver cell types in the multiome data. The inner-most track represents links between the ligand receptor pairs, with the thickness of edge indicating log2FC of strength of interaction between crypto and healthy conditions and the colour indicating upregulated (red) or downregulated (blue) LRIs between crypto and healthy conditions. (right) Heatmap of the top 5 up- and downregulated ligand–receptor interactions (LRIs) between crypto and healthy conditions, across sender cell types for each receiver cell type. The LRIs are shown on the y-axis, sender cell types are shown on the x-axis, with separate facets corresponding to receiver cell types. Ligands and receptors discussed in the text are indicated in bold.

Because this disease heterogeneity was reflected by the differential expression of multiple transcription factors from our high-confidence network (Fig. 5D, S8F, G), we tested whether our SCENIC+ eRegulons accurately predicted the downstream consequences of these TF perturbations. To quantify this, we assessed the concordance between every TF-cluster pairs involving TFs from positive eRegulons. Across three patients, between 32.8% and 62.5% of these instances demonstrated significant directional concordance between the expression shifts of a perturbed TF and its downstream target network (Fisher’s exact test, FDR < 0.05). To visualize these specific regulatory dynamics, we normalized target gene expression against cluster-average shifts. This revealed clear concordant and inverse relationships for positive and negative eRegulons, respectively (Fig. 5E-H, S8H, I). This *in vivo* directional concordance underpins the predictive value of our computational framework, providing orthogonal support for the inferred gene regulatory network. Consequently, the network correctly identified candidate downstream dysregulation of critical spermatogenic regulators such as NR3C1 ^80^ and DMRTB1, a PIWIL-pathway regulator ^81^, in spermatogonia (Fig. 5E, G).

Because functional validation of these dysregulated target genes is not feasible in humans, we utilized the International Mouse Phenotyping Consortium (IMPC) database as an orthogonal reference for gene essentiality in male fertility. By cross-referencing our data against established murine knockout models, we sought to determine whether the specific network perturbations observed in our cohort converge on genes strictly required for mammalian spermatogenesis. The dysregulated network in Crypto 2 was highly enriched for orthologues whose ablation causes male sterility in mice (Fisher’s Exact Test p = 0.003). These targets encompass critical structural and meiotic regulators, including the acrosomal protease inhibitor SPINK2 ^82^, the flagellar protein CCDC34 ^83^, and the synaptonemal complex associated protein HORMAD1 ^84^ (Fig. 5I).

Finally, we assessed how these intracellular transcriptional shifts impact the spatial cellular niche using our cell-cell communication (CCC) framework. Global analysis revealed a severe reduction in intercellular signalling in the disease state (Fig. 5J, S8J, S9A, B). Within the spermatogonial niche (cluster 5) of Crypto 2, critical maintenance signalling pathways were attenuated, including GRM7 ^85^, PTN ^86,87^, and PENK ^88^. Similarly, the somatic support architecture exhibited profound collapse: Sertoli cells (cluster 24) lost crucial germ-cell adhesion and developmental signals (CADM1 ^89^, NRG1 ^90^, and ERBB4 ^91,92^), while PTMs (cluster 8) exhibited a compensatory increase in EGFR-associated signalling, indicative of a severe tubular remodelling response ^93^.

Collectively, these *in vivo* perturbation data support the predictive value of our multiomic network. By seamlessly integrating spatial architecture, chromatin accessibility, and transcriptional regulation, this atlas provides a same-nucleus, spatially-anchored framework for resolving the heterogeneous molecular pathologies driving human male infertility.

## Discussion

Here, we characterize the dynamics of chromatin accessibility, gene expression, and spatial architecture throughout human spermatogenesis, utilizing this data to functionally annotate the molecular pathology of male infertility. While previous studies have independently analyzed scRNA-seq and scATAC-seq in the human testis ^37–39^, the reliance on probabilistic label-transfer limited the resolution of regulatory networks. By utilizing single-nucleus multiomics to capture both modalities within the same cells, we observed that promoter accessibility frequently persists throughout spermatogenesis despite cluster-specific gene expression. This disconnect underscores the limitations of inferring transcription from promoter accessibility alone. By bypassing this limitation, we inferred a robust gene regulatory network comprising 62 high-confidence eRegulons driven by known stage-specific regulators (e.g., DMRT1) and novel candidate regulators (e.g., KLF17). Compared to traditional transcript-only network inference, our stringent criteria for predictive cis-regulatory mapping naturally filters out co-expressed transcripts that lack direct chromatin-level evidence, highlighting the enhanced resolution of true multiomic profiling. Crucially, by inferring networks from same-nucleus matched modalities, we resolve the developmental specificities of these high-confidence regulators and experimentally validate the spatial localization of key drivers, namely, DMRT1, ZBTB43, CREM and EGR1, in situ at the protein level.

Analysis of this regulatory network revealed a distinct topological architecture: regulators active in spermatids occupy the network periphery, exhibiting lower betweenness centrality than regulators maintaining spermatogonia and somatic cells. Previous research correlates network centrality with evolutionary rates, wherein younger, faster-evolving genes exhibit lower centrality ^95,96^. The peripheral localization of spermatid transcription factors is therefore highly concordant with recent findings that genes expressed in late spermatogenesis are younger and evolving more rapidly than those expressed during early differentiation ^23^. This topology likely provides a permissive framework for rapid evolutionary adaptation in post-meiotic cells without disrupting the highly conserved regulatory core governing the basal niche ^97,98^. As mice remain the primary *in vivo* model for reproductive genetics, this rapid evolutionary divergence highlights a potential translational challenge. We anticipate that this human eRegulon atlas will serve as a valuable comparative reference to evaluate the evolutionary conservation of specific TF-enhancer modules, helping to contextualize findings from murine models and resolve human-specific regulation.

The physiological relevance of these regulatory modules is further supported by mapping them to their native spatial context. By integrating our single-nucleus multiome data with spatial transcriptomics, we anchored specific structural and immunological effectors to discrete intercellular signalling axes. Within the germline, our spatially-aware cell-cell communication analysis localized the receptors for FGF12 and NLGN1—transcripts implicated in idiopathic infertility ^99^ and spermatogonial stem cell maintenance ^100^ respectively—directly to the spermatogonial niche. In the somatic compartment, the spatial neighbourhood analysis resolved extensive CADM1 homotypic interactions between Sertoli and germ cells ^89^. This directly supports recent observations of robust *CADM1* expression in the Sertoli niche ^23^ and provides a refined human framework for understanding the profound spermatogenic failure observed in murine Cadm1 knockout models ^89^. Furthermore, interactions involving DCN ^101^ and PECAM1 ^102,103^ mapped strictly to extratubular somatic and immune compartments, linking genes commonly dysregulated in non-obstructive azoospermia to specific structural and inflammatory cell populations.

This integrated regulatory map provides a useful framework for interpreting non-coding genetic variation. In contrast to other complex traits, the genetic architecture of male infertility remains poorly defined. Current infertility GWAS are underpowered, yielding loci with unclear mechanisms ^41^. Consistent with this, while our partitioned heritability (S-LDSC) analysis revealed positive coefficient estimates across meiotic cell types, these signals did not survive multiple-testing correction, directly reflecting the limited statistical power of available cohorts. Because cell-type-specific regulatory elements are difficult to isolate from heterogeneous testis tissue, the functional interpretation of non-coding variants has stalled. Here, we maximized the utility of existing GWAS data by projecting statistical associations onto our deterministic network. We mapped several non-coding SNVs to clinically relevant target genes, including *TNFRSF14* ^68^ and *SORBS1* ^69^ in the peritubular myoid niche, *MMEL1* ^66,67^ in spermatids and *APOL2* in endothelial cells ^73^. Additionally, cell-type-resolved mapping can identify candidate regulatory elements in non-germline populations for genes with established germline functions. For example, while SYNJ2 is best characterized for its structural role in germ cells ^74,75^, the rs74863147 variant overlaps an enhancer accessible in endothelial cells. Whether this region exerts a functional regulatory effect in the testis remains to be established. By pinpointing the specific transcription factors and cell types through which these variants operate, this atlas moves correlative genetic associations towards testable mechanistic hypotheses.

While our multiomic pipeline identifies thousands of candidate distal regulatory elements, functionally validating these specific sequences highlights a major bottleneck in the field. As demonstrated by our comparison with recently published in vitro-validated enhancers ^25^, isolating bona fide distal elements outside their native biological context remains highly challenging. Although our *in vivo* atlas successfully captures established functional loci like *PRDM9*, the lack of broader concordance underscores that current *in vitro* enhancer reporter assays often lack the robust cellular context and negative controls necessary to accurately recapitulate the highly specialized, dynamic microenvironment of the adult testis. Consequently, while developing high-fidelity *in vitro* reporter systems remains a critical objective, we currently rely on the integration of established genetic variants and clinical perturbation models to functionally anchor this regulatory network.

Finally, by profiling clinical cryptozoospermia samples, we provide *in vivo* support for the predictive value of this network. Despite sharing a severe depletion of spermatids, the three infertile individuals exhibited distinct molecular etiologies, underscoring the extreme phenotypic heterogeneity of spermatogenic failure. While this vast heterogeneity precludes defining a singular etiology for cryptozoospermia, these distinct molecular profiles provided an opportunity to use the patient samples as individualized *in vivo* perturbation models.

Crucially, when transcription factors were disrupted in these patients, their predicted target genes exhibited concordant dysregulation. In Crypto Rep2, this dysregulated downstream network was significantly enriched for essential fertility genes whose ablation causes sterility in mice, including SPINK2 ^82^, HORMAD1 ^104^, and FSIP2 ^105–107^. Furthermore, the atlas captured the consequent breakdown of the somatic support niche, identifying the specific attenuation of spermatogonial maintenance signals (PTN) and Sertoli cell adhesion pathways (CADM1, ERBB4). Rather than merely cataloging differential expression, this network highlights candidate regulatory nodes associated with niche collapse.

Beyond providing a descriptive atlas of human spermatogenesis, this study illustrates several features that distinguish regulatory multiomic maps of the testis from analogous atlases in other organs. First, the human testis is a highly structured developmental tissue in which germ-cell differentiation unfolds within a spatially organized somatic niche, requiring regulatory inference to be interpreted in the context of both cell state and tissue architecture. Second, by integrating single-nucleus multiomics with spatial transcriptomics, we were able not only to define cell-type-specific eRegulons, but also to place these programmes within localized somatic–germline signalling environments of the seminiferous tubule. Third, we utilized rare clinical biopsies as in vivo perturbation models to assess whether the inferred regulatory framework could capture the altered molecular states and the secondary impacts on the somatic niche associated with spermatogenic failure. Together, these features move the study beyond a static reference atlas towards a spatially anchored and perturbation-informed framework for interpreting human developmental regulation, infertility-associated genetic variation, and clinically heterogeneous testicular pathologies.

Several limitations of this study should be considered. First, compared with other complex traits, currently available male infertility GWAS remain relatively underpowered, limiting the number of non-coding risk loci that can currently be interpreted in a regulatory framework. Second, certain somatic populations (such as Leydig and Sertoli cells) are underrepresented in our baseline dataset. As noted in previous testis atlases ^16,108–110^, this is a common technical limitation primarily driven by the high relative abundance of post-meiotic germ cells, which effectively dilutes rare interstitial populations in unsorted tissue suspensions. Indeed, the severe depletion of spermatids in our cryptozoospermia samples removed this unmasking effect, enabling robust identification of these somatic populations. However, deeper profiling of rare extratubular cells in healthy tissues will ultimately require targeted sorting or enrichment strategies. Third, our spatial cell-cell communication framework prioritizes interactions between physically adjacent cells, which robustly captures contact-dependent and local paracrine signalling but may underrepresent the influence of long-range, soluble endocrine factors operating within the testis. Finally, while our clinical multiomic data provide an opportunity to examine regulatory perturbation *in vivo*, the number of patient samples remains constrained by the availability of fresh clinical biopsies. Larger cohorts spanning diverse infertility subtypes will be required to capture the full molecular landscape of male infertility and further refine these inferred disease-associated regulatory programmes.

In summary, we provide a same-nucleus, spatially-resolved atlas of the gene regulatory networks governing human spermatogenesis. This resource helps address a major gap in reproductive genomics by providing the necessary regulatory context to interpret non-coding variation and heterogeneous clinical phenotypes. Ultimately, we envisage that this functional map may improve interpretation of whole-genome sequencing in infertile men, inform future therapeutic strategies, and provide molecular benchmarks for advancing *in vitro* gametogenesis.

## Methods

### Selection of human testicular biopsies

We obtained testicular biopsies of patients from the Centre of Reproductive Medicine and Andrology (CeRA) in Münster, Germany. Normal spermatogenesis samples were obtained from patients who had been diagnosed with obstructive azoospermia due to a previous vasectomy, a congenital malformation or physical obstruction of the vas deferens, but had normal testicular volume, normal FSH levels and qualitatively and quantitatively normal spermatogenesis. Cryptozoospermic patients had a sperm concentration < 0.1 million/ml (cryptozoospermia) in the ejaculate, a reduced testicular size, and, in most cases, elevated FSH levels due to a severe impairment of spermatogenesis itself. All patients had spermatozoa in their TESE samples, regardless of histological results. A complete list of the measured clinical parameters can be found in Supplementary Tables S1 and S4 for normal and cryptozoomspermic patients respectively. When undergoing surgery for microdissection testicular sperm extraction (mTESE) or histological evaluation at the Department of Andrology (University Hospital in Münster, Germany), one additional testicular sample was excised for this study after written informed consent. Ethical approval was obtained for this study (Ethics Committee of the Medical Association of Westphalia-Lippe and University of Münster 2010-578-f-S).

### Preparation of single cell suspensions from human adult testis biopsy

A two-step enzymatic digestion was performed as detailed in ^111–113^. Briefly, the tissue sample was placed into 6 cm culture dish with MEM-alpha media (Life Technologies GmbH, Gibco, Darmstadt, Germany) and dissected into fragments of 1mm or less using a blade. It was then transferred into a 15 ml tube and centrifuged for 5 minutes at 1500 rpm and 10°C. The cell pellet was resuspended in MEM-alpha media containing 1mg/ml filter sterilized collagenase IA (C9891, Sigma) and incubated at 37°C until tubules separated. The tube was inverted every 5 minutes. The reaction was stopped by adding 5 ml MEM-alpha media with 10% FBS and 1% Pen/Strep. It was centrifuged at 10°C at 1500 rpm for 4 minutes, the supernatant discarded and incubated with 2ml filter sterilised Hank’s balanced salt solution (Life Technologies GmbH, Gibco) containing 2 mg Trypsin (27250-018, Gibco) and 11 mg DNase I (DN25, Sigma) for 7-10 minutes at 37°C. The tube was inverted every 3 minutes.

This was followed by strong pipetting to obtain a single-cell suspension. The reaction was stopped as outlined above and cells were washed twice with MEM-alpha. Then, erythrocytes were removed by incubating the cells with haemolysis buffer (0.83% NH4Cl solution) for 3 minutes at room temperature. Enzymatic digestion was stopped as outlined above. The cells suspension was filtered through a 70 μm strainer to remove cell clumps and debris. It was pelleted, washed and the cells were resuspended in 2 ml medium prior to cell counting using the trypan blue exclusion method.

### Generation of paired scATAC-seq and scRNA-seq libraries

200,000-400,000 cells were centrifuged at 500g for 10 minutes at 4°C. The supernatant was discarded, and the pellet resuspended in 50 μl Wash Buffer (20 mM HEPES pH 7.5, 150 mM NaCl, 0.5 mM Spermidine, 1X Protease Inhibitor Cocktail). It was centrifuged at 500g for 10 minutes at 4°C and supernatant was discarded. The pellet was incubated with 45 μl Lysis Buffer (wash buffer with 0.01% NP40, 0.01% Digitonin, 1mM DTT and 1 U/μl RNAse inhibitor (3335399001, Sigma)) for 25-30 seconds on ice. The reaction was stopped with 50 μl Neutralising Buffer (lysis buffer with 2mM EDTA). It was again centrifuged at 500g for 10 minutes at 4°C, the supernatant discarded, and the pellet washed with 45 μl 1X Dilute Nuclei Buffer (20X Nuclei Buffer provided with 10x scATAC + GEX multiome kit). The pellet was resuspended in 10 μl 1X Dilute Nuclei Buffer solution and 2 μl sample was taken for counting with Trypan Blue.

9000-10,000 nuclei were loaded for the 10x scATAC + GEX multiome transposase reaction and the library prepared according to manufacturer’s instructions in Chromium Next GEM Single Cell Multiome ATAC + Gene Expression Reagent Kits User Guide (Revision A). The libraries were sequenced in NovaSeq 6000 with paired end sequencing.

RNA-seq libraries were sequenced to a depth of 40,000 read pairs/nucleus and ATAC-seq libraries to a depth of 50,000 read pairs/nucleus according to 10x sequencing guidelines.

### scATAC-seq and scRNA-seq analysis

#### Read mapping and quantification

The fastq files of individual samples were processed with cellranger-arc v2.0.0 and mapped to refdata-cellranger-arc-GRCh38-2020-A-2.0.0.tar.gz. For each sample, cellranger-arc count was used with default parameters to generate a filtered peaks x cells counts matrix and a filtered genes x cells counts matrix. The filtered matrices of each sample were loaded into R and further processed with Seurat v4.1.1^43^ and Signac v1.7.0 ^114^.

#### RNA library QC

Ambient RNA contamination was estimated and removed using soupX v1.6.1 ^115^. Next, doublets were removed using scDblFinder v1.10.0 ^116^, using a doublet rate of 0.077 or 0.069, corresponding to the multiplet rate for 10,000 and 9,000 recovered cells respectively in the 10x multiome scATAC + GEX user guide (Revision A). Cells with > 20% mitochondrial reads, fewer than 500 UMIs or fewer than 250 detected genes were removed. These cut-offs were determined using the isOutlier() function from scater v1.24.0 ^117^ or by visually examining the distribution with the aim of retaining as many high quality cells as possible.

#### ATAC library QC

Doublets were removed with using scDblFinder v1.10.0 ^116^, using a doublet rate of 0.077 and 0.069 for 10,000 and 9,000 recovered cells respectively, and aggregateFeatures = TRUE. Cells with fewer than 150 detected peaks or fewer than 250 fragments were removed. These cut-offs were chosen as described above. Cells with FRiP < 0.20 or TSS enrichment < 2 were removed.

#### Peak calling

Peaks were called on the remaining high quality cells grouped by Cellranger-identified cluster, using macs2 ^118^ via the Seurat function CallPeaks(). Peaks on non-standard chromosomes and those in genomic blacklist regions were removed. A new assay was created by quantifying counts in each of the remaining peaks with function FeatureMatrix() from Seurat taken for all downstream processing.

### Normalization, sample integration, dimensional reduction

The RNA and ATAC-seq assays of the biological replicates were separately integrated using Seurat.

#### RNA-seq assay

Genes x cells matrices were normalised with scran v1.24.0 ^119^ as follows. The Seurat object was converted to a SingleCellExperiment (SCE) object and clustered using the quickCluster() function. Size factors were calculated using computeSumFactors(). The logNormCounts() function from scuttle v1.6.2 ^117^ was used to calculate normalised counts with log = FALSE. The normalised counts were then log-transformed with the log1p() function and added to the data slot of the Seurat object.

The top 4000 variable features were identified with Seurat’s FindVariableFeatures() function with selection.method = "vst". The datasets were then integrated using the Seurat functions SelectIntegrationFeatures(), FindIntegrationAnchors(), and IntegrateData(). Finally, the combined integrated matrix was then scaled and centred with function ScaleData() from Seurat and PCA was performed using RunPCA().

#### ATAC-seq assay

The union of peaks called in each individual snATAC-seq dataset was taken and all datasets were re-quantified using this unified peak set. The bottom 5% of features were removed and TF-IDF and SVD were carried out to produce an LSI matrix. Datasets were then merged and the LSI calculation repeated. The LSI embeddings were integrated using the functions FindIntegrationAnchors() (using all peaks and reciprocal LSI) and IntegrateEmbeddings().

#### Weighted nearest neighbour (WNN) identification, clustering, and cluster annotation

The Seurat function FindMultiModalNeighbors() was used on the integrated RNA data (PCA dimensions 1:20) and ATAC data (LSI dimensions 2:20) with k.nn = 25 for normal samples and k.nn = 30 when combining normal and crypto samples. Clustering was carried out on the resulting WNN network using the FindClusters() function with res = 0.8 for normal samples and res = 0.9 when combining normal and crypto samples. Clusters were annotated based on the expression of key marker genes ^26,44^.

### scRNAseq analysis for scRNA vs snRNA comparison

#### Read mapping

The fastq files of individual samples were processed with cellranger v6.0.1 and mapped to refdata-gex-GRCh38-2020-A.tar.gz. For each sample, cellranger count was used with default parameters to generate a filtered genes x cells counts matrix.

#### Quality control filtering

The filtered matrices of each sample were loaded into R and further processed with Seurat v4.0.3. Doublets were removed using scDblFinder() v1.10.0 with the following doublet rates: 0.008 for Guo et al. 2018 ^16^, 0.046 for Di Persio and Tekath et al. 2021 ^26^, and 0.077 or 0.069 for 10,000 or 9,000 cells recovered in this study, according to the respective 10x user guides. Cells with > 24% mitochondrial reads, nFeature_RNA < 350 (< 1000 for this study) and > 9500 were removed. These cut-offs were based on function isOutlier() from scater v1.20.1 or by visually examining the distribution with the aim of retaining as many high-quality cells as possible. Sample N1 from Di Persio and Tekath et al. 2021 was excluded from the analysis due to high ambient RNA contamination that resulted in higher than estimated number of droplets being called as cells by Cellranger.

Each dataset was normalised with scran v1.24.0 as described above for multiome data. The top 4000 variable features were identified with Seurat function FindVariableFeatures() with selection.method = "vst".

#### Normalization, sample integration, dimensional reduction

Datasets were then integrated with Seurat using the Seurat functions SelectIntegrationFeatures(), FindIntegrationAnchors(), and IntegrateData(), with normalization.method = "LogNormalize”. The integrated assay was scaled and centred with Seurat function ScaleData() with all genes as features and PCA was performed using RunPCA().

#### Clustering and cluster annotation

Cells were clustered using Seurat function FindClusters() on the integrated RNA with dims 1:30 and resolution = 0.5. Clusters were annotated based on key marker gene expression, as used in Di Persio & Tekath et al. 2021 ^26^.

### Trajectory inference

Trajectory inference was carried out using Slingshot v2.4.0 ^45^, using the first 20 dimensions of the PCA reduction of the RNA data after ambient RNA removal. Only germline clusters were included. The cluster corresponding to Spermatogonia State 0 was specified as a trajectory start point. This produced two lineages. The main lineage shown in Fig.1C (inset) includes > 99% of germline cells. A secondary lineage only includes clusters representing undifferentiated spermatogonia (37% of germline cells, not shown).

### Differential gene expression

Identification of cluster-specific enriched genes was carried out using the FindAllMarkers() function from Seurat, using method “MAST” ^120^ with biological replicate as a latent variable. Only positively enriched markers with an adjusted p value < 0.05 were kept.

Identification of differentially expressed genes between normal and cryptozoospermic samples was carried out for each cluster independently using Seurat FindMarkers() with method “wilcox”. Genes with an adjusted p value < 0.05 and an absolute log2 foldchange in expression of > 1 were kept.

### GO term enrichment

The top 100 most significantly enriched genes per cluster were used as input for GO term enrichment analysis with clusterProfiler (v4.7.1.003) function enrichGO ^121^, using all genes included in the differential gene expression analysis as the background gene set. The results were simplified to remove terms with high semantic similarity using method = “Wang” and cutoff = “0.7”. For visualisation, the most significantly enriched (based on BH-adjusted p value) term per cluster was selected and enrichment of these terms across all clusters plotted. For GO term enrichment in cluster-specific differentially expressed genes between normal and Crypto samples, all genes with BH-adjusted p value < 0.05 and an absolute log2 foldchange in expression of > 1 were selected. Up- and downregulated genes were analysed separately, as described above. The results are supplied in Supplementary Table S9.

### Differential accessibility

Identification of differentially accessible peaks between normal and cryptozoospermic samples was carried out for each cluster independently using Seurat FindMarkers() with method “LR”. The total number of reads in peaks was used as a latent variable and only peaks with signal in at least 5% of cells in the cluster were assessed. Peaks with an adjusted p value < 0.05 were kept.

### Motif activity

Motif activity was calculated using the Arch v1.0.2 R ^122^ implementation of ChromVar ^123^. Cellranger output filtered matrices were pre-processed using ArchR, keeping cells with a minimum of 80 fragments and a minimum TSS enrichment value of 1.45. Cells were then further filtered to keep only those that had passed the filtering and quality control procedures described above. Consensus motifs from Vierstra et al. 2020 ^49^ obtained from https://resources.altius.org/~jvierstra/projects/motif-clustering-v2.0beta/ were used for motif activity calculations using ArchR with default parameters. The resulting ChromVar Z-scores were then added back to the integrated Seurat object for downstream analysis.

### Generation of spatial transcriptome library

Formalin fixed paraffin embedded human testicular biopsies were obtained from the Imperial College Healthcare Tissue Bank (ICHTB) collection under REC:22/WA/0025, HTA license 12275 and project license R19000. Samples were collected from a 37-year-old individual who underwent orchiectomy for suspected gonadal tumours. The tissues were sectioned in 5µm slices and mounted onto Xenium slides according to 10X Genomics workflow CG000578. Histological assessment was performed using DAPI and H&E stains. Spatial transcriptomic profiling was performed in collaboration with the Genomics Facility at the MRC Laboratory of Medical Sciences. Briefly, sections were deparaffinized and decrosslinked following workflow CG000580, then underwent probe hybridisation for 17 h at 50°C with a Xenium standalone custom gene panel, followed by post-hybridisation wash, ligation, amplification, autofluorescence quenching, and nuclear staining per workflow CG000582. Finally, the slides are loaded onto Xenium Analyzer for barcode decoding and imaging following workflow CG000584 using Xenium onboard analysis (v1.9.0.0).

### Tubule segmentation in spatial transcriptome

Post Xenium, the quencher was removed, and the slides were stained with H&E according to protocol CG000613. Seminiferous tubules were segmented from H&E-stained Xenium tissue sections using the QuPath extension Segment Anything Model (v0.5.0) ^124–126^ implemented in QuPath (v0.4.4) ^127^. Segmented tubule boundaries were exported as polygon annotations and transformed into the Xenium coordinate space using a transformation matrix derived from key-point alignment on Xenium Explorer (v.3.2.0) ^128^.

Transformed tubule polygons were intersected with Xenium cell segmentation geometries to assign individual cell identifiers to seminiferous tubules.

### Immunofluorescence staining

Human testis tissue sections were cut at 5 μm thickness using a Leica CM3050S cryostat. Antigen retrieval was performed by boiling slides in 10 mM sodium citrate buffer, pH 6.0–7.0, for 10 min. Sections were blocked in PBS containing 0.01% Triton X-100 and 10% donkey serum (Merck, D9663) for 2 hours at room temperature. Slides were incubated overnight at 4 °C in a humidified chamber with primary antibodies (OCT4 ab181557) diluted in blocking buffer. After washing in PBS, sections were incubated with Alexa Fluor 594-conjugated donkey anti-rabbit secondary antibody (Thermo Fisher Scientific; 1:300) for 1 h at room temperature. Nuclei were counterstained with DAPI, and sections were mounted using VectaShield antifade mounting medium (Vector Laboratories, H-1000). Images were acquired on a Leica SP5 and processed in Fiji (ImageJ2, 2.9.0/1.53u).

### Pre-processing of Xenium spatial data

#### QC filtering, Normalisation, dimensional reduction

Xenium data was processed on-machine using Xenium Ranger v1.9.0.0, then re-segmented using Xenium Ranger v2.0.0.12, to benefit from its improved segmentation algorithm.

Normalisation and dimensionality reduction were carried out using Seurat (v5.0.3).

First, cells with zero counts were removed. Expression data was normalised using SCTransform, PCA was performed, and UMAP transformation was carried out using the first 20 PCs.

#### Clustering and cluster annotation

Clustering was carried out using Xenium Ranger (v2.0.0.12). Clusters were manually annotated based on a combination of approaches: marker gene expression, identification of cluster-specific upregulated genes, and label transfer using RCTD from the spacexr package (v2.2.1) ^129^.

The same set of marker genes used for the sn-multiome analysis was applied here. These markers were derived from Di Persio & Tekath et al. (2021) ^26^ and served as the primary basis for cell type annotation. Cluster-specific upregulated genes were identified using FindAllMarkers() from Seurat (v5.0.3). To aid in cluster annotation, we visualised expression of these upregulated genes across both the Xenium clusters and the sn-multiome clusters. Label transfer from sn-multiome data to the Xenium data was carried out using RCTD from the spacexr package (v2.2.1). The reference dataset was the gene expression data from the merged normal sn-multiome samples, after ambient RNA correction. “Unclear” clusters were removed. RCTD was run with doublet mode “doublet”, to return a max of two possible labels.

### Tubule type classification

Post tubule segmentation based on H&E staining, cells were assigned to tubules using custom python scripts. First, Xenium data was converted to SpatialData Zarr format using spatialdata v0.2.2 ^130^ and the Xenium reader from spatialdata-io v0.1.4, slightly modified to allow reading of re-segmented data. The shapely package (v2.0.5) ^131^ was used to assign cells to tubules by taking the cell centroid (central point of cell circles element of the SpatialData object) and identifying the tubule containing this point, if any, using the shapely.contains function. Tubules were characterised by creating a matrix of the proportion of each cell type cluster within each tubule (i.e. each row is a tubule, each column is a cell type cluster). The distance matrix between rows was calculated using the built-in dist function in R (v4.2.0). This distance matrix was clustered using the built-in hclust function with method “ward.D”. The cutree function was used to identify two groups of tubules.

### Spatial trajectory inference

Genes detected in fewer than 10 cells were excluded. Gene expression counts were normalised to a total of 1 × 10⁴ per cell and log1p transformed. A region of interest was manually defined, and the dataset subset for downstream analysis. Spatial trajectory inference was performed using SpaTrack (v0.1.1) ^54^. Firstly, cluster 2: Spermatogonia was specified as the root population. The weightage of gene expression information and spatial location information was calculated using calc_alpha_by_moransI() and the resulting values of alpha1= 0.845 for gene expression and alpha2=0.155 for spatial information was used to compute the transition probabilities between cells and using the top 50 principal components. Pseudotime was calculated from the defined root cells and velocity vectors estimated using 50 neighbours based on cell positions (and no gene expression-based neighbours), with a smoothing factor of 0.5 and grid density of 1.0.

### Neighbourhood enrichment analysis

Neighbourhood enrichment z-scores were calculated using Squidpy (v1.5.0) ^132^. Cells with zero transcript counts were excluded from analysis. A spatial graph was constructed using the generic coordinate system and Delaunay triangulation. Neighbourhood enrichment was then calculated with 1,000 permutations to assess statistical significance.

### Cell-cell communication analysis

Ligand-receptor interactions were inferred using scDiffCom (v1.0.0) ^55^ with full detection and differential analysis for human data. Genes that are expressed in at least 20 cells and in ≥20% of cells within a given group were retained, and cell types with less than 20 cells were removed from the analysis. Statistical significance was assessed using 10,000 permutation iterations. CCIs with scores in the lowest 20% or with a Benjamini–Hochberg–adjusted specificity p-value > 0.01 were excluded from detection.

For circos plot visualization, only ligand–receptor interactions with positive scores from Squidpy neighborhood enrichment analysis were retained. As meiotic cells were not resolved as a distinct population in the spatial dataset, interactions involving these cells were excluded from the final set of LRIs selected for visualisation from the multiome data.

Differential LRI analysis between Normal and Crypto samples was performed using the same workflow, with condition1 specified as Normal and condition2 as Crypto for each pairwise comparison.

### SCENIC+

ATAC-seq counts and cell metadata were exported to text files from the integrated Seurat object and used to construct a cisTopic object using pycisTopic v1.0.2.dev8+g848f78b. Topic modelling was carried out using pycisTopic with 500 iterations, alpha = 50, eta = 0.1, and numbers of topics between 5 and 40. The optimal number of topics was determined to be 30 based on the suggested metrics. This selected set of topics was then binarized in two ways: using the Otsu method and by taking the top 3000 regions per topic. In addition, differentially accessible regions across clusters were identified using default parameters. These sets of regions were then used as input into pycisTarget (v1.0.2.dev8+g48af509). pycisTarget was used to calculate motif enrichment using the provided motif rankings and scores for motifs from the 2022 SCENIC+ motif collection (database version “hg38_screen_v10_clust”) ^13^.

The cisTopic object and motif enrichment results were combined with and RNA-seq counts after ambient RNA removal exported from Seurat to construct a SCENIC+ object ^13^. SCENIC+ (v0.1.dev446+g5cf2469) was then run with default parameters. Search space for putative regulatory regions was 150kb upstream and downstream of each gene. Standard filtering was applied and returned 521 eRegulons, with a total of 471 TFs and 13,685 target genes. Per-cell enrichment (AUC) of eRegulon genes and regions was calculated using the AUCell function. High confidence eRegulons were selected as those where the absolute correlation between TF expression and gene-based AUC was more than 0.6, the absolute correlation between TF expression and region-based AUC was more than 0.6, and the correlation between gene-based and region-based AUC was more than 0.4.

### Network analysis

The high-confidence eRegulons were used to construct a network where nodes represent TFs and target genes. The network was constructed using tidygraph (v1.2.2) ^133^ and visualised using ggraph (v2.1.0) ^134^. Centrality metrics were calculated using tidygraph.

### TF footprinting

In order to carry out TF footprinting for each cluster, the subset-bam tool from 10x (v1.1.0) was used to produce bam files containing reads for cells assigned to each cluster. Individual replicates were merged using samtools (v1.16.1) ^135,136^ to produce one bam file per cluster. These bam files were used as input to TOBIAS (v0.13.3) ^57^ for footprinting of motifs from Vierstra et al. 2020 ^49^ obtained from https://resources.altius.org/~jvierstra/projects/motif-clustering-v2.0beta/ within peaks identified in any cluster from the Seurat object. Regions falling into the blacklist regions included in the Signac package were excluded. TOBIAS was run in time-series mode on germline clusters only.

TOBIAS PlotAggregate was used to aggregate TF binding scores in 120bp windows across all detected motifs within peaks. For each motif and cluster, the average signal in the central 20bp around the motif and average signal in the flanking regions were calculated. Motifs with a notable footprint have less signal in the central region than in the flanking regions (a positive footprint depth); motifs lacking footprint signal in all clusters were removed from analysis, leaving 296 motifs of interest.

### DMRT1 ChIP-seq analysis

DMRT1 ChIP-seq data was obtained as fastq files from the authors ^56^. Reads were aligned using bowtie2 (v2.5.0), sorted and filtered using samtools (v1.16.1) to keep only primary alignments with mapping quality at least 30. Duplicates were marked using sambamba markdup (v0.8.1). Replicates were merged using samtools (v1.16.1). Coverage tracks for individual and merged replicates were generated using bamCoverage (deeptools v3.5.1), with bin size 10 bp, CPM normalisation, extending alignments to 200 bp, and ignoring duplicate alignments and those with mapping quality < 30. Peaks were called for individual and merged replicates using macs2 (v2.2.9.1) with a fixed alignment extension to 200 bp and a q-value cutoff of 0.2.

The regioneR package (v1.28.0) ^137^ was used to test for significance of overlap of DMRT1 ChIP-seq peaks and DMRT1 eRegulon target regions compared to all other SCENIC+ eRegulon target regions, using resampleRegions with default parameters and 1000 permutations.

### Immunohistochemistry for TF validation

Testicular biopsies (n = 3) were dissected, placed in histology cassettes, and fixed overnight in Bouin’s solution. The tissues were subsequently transferred to 70% ethanol, dehydrated through a graded ethanol series, and embedded in paraffin. Immunohistochemical staining was performed on 5μm paraffin sections. Sections were dewaxed with AppiClear (Applichem, A4632.2500), rehydrated through a descending ethanol series, and rinsed in distilled water. Heat-induced antigen retrieval was performed using sodium citrate buffer (pH 6.0). Endogenous peroxidase activity was blocked with 3% hydrogen peroxide (Hedinger, GH06708-001), followed by blocking of non-specific primary and secondary antibody binding with 5% bovine serum albumin (BSA; Sigma-Aldrich, A9647) and 25% goat serum (Sigma-Aldrich, G6767-100ML), respectively. Sections were incubated overnight at 4°C with the following primary antibodies: rabbit anti-ZBTB43 (Sigma-Aldrich, HPA016825, 1:10), rabbit anti-CREM (Sigma-Aldrich, HPA001818, 1:100), mouse anti-DMRT1 (Santa Cruz, sc-377167, 1:100), and rabbit anti-EGR1 (Sigma-Aldrich, HPA029937, 1:50). For negative controls, one section per patient was incubated with rabbit (Sigma-Aldrich, I5006) or mouse (Sigma-Aldrich, I5381) IgG instead of the corresponding primary antibody. The following day, sections were incubated with species-specific biotinylated goat secondary antibodies (anti-mouse, Abcam, ab5886; anti-rabbit, Abcam, ab6012; both 1:100), followed by streptavidin–horseradish peroxidase (Sigma-Aldrich, S5512, 1:500). Peroxidase activity was visualized using 3,3′-diaminobenzidine tetrahydrochloride (DAB; Sigma-Aldrich, D5905), and the reaction was terminated by rinsing in distilled water. Nuclei were counterstained with Mayer’s hematoxylin (Sigma-Aldrich, 1.092.490.500). Finally, the slides were dehydrated through an ascending ethanol series, cleared with AppiClear, and mounted with Merckoglas (Sigma-Aldrich, 1.039.730.001). Whole-slide images were acquired using an Evident Slideview VS200 Universal whole-slide scanner and analyzed with QuPath software.

### GWAS data integration

rsIDs for SNVs associated with infertility in published GWAS studies were obtained from the NHGRI-EBI GWAS catalog ^59,62,138^ and directly from ^41,60,61^. The LDLinkR package (v1.4.0) ^63^ was used to obtain lists of all SNVs within 500kb of the lead SNVs and in linkage disequilibrium with them in the respective study population (Han Chinese or European ancestry or all ancestry)^63,64^, using genome build “grch38_high_coverage”. SNVs with r^2^ > 0.8 were selected for further analysis. Prior to assessing SNV–regulatory region overlaps, SCENIC+ regions overlapping gene promoters were excluded.

### LDSC partitioned heritability

Partitioned heritability was estimated by stratified LD score regression using LDSC v1.0.1 ^139^ applied to GWAS summary statistics ^140^. Pre-computed LD scores, regression weights and allele frequencies derived from 1000 Genomes Project Phase 3 European-ancestry reference panels were obtained from the Broad Institute LDSC repository (https://storage.googleapis.com/broad-alkesgroup-public/LDSCORE/). The major histocompatibility complex locus was excluded from all analyses owing to its atypical linkage disequilibrium architecture. Analyses covered 1,093,784 HapMap3 variants and were conditioned on the 97-annotation baseline model (v2.2).

Cell-type-specific heritability was estimated across 21 cell types from adult human testes using MACS2 peaks called from snATAC-seq data, with each annotation added individually to the baseline model. Significance was assessed on the regression coefficient τc, a one-sided test of whether an annotation contributes to trait heritability conditional on the baseline annotations.

To contextualise findings, we analysed European GWAS summary statistics for female infertility and other phenotypes with established relevance to male reproductive health: total testosterone ^141^, systemic lupus erythematosus ^142^, rheumatoid arthritis ^143^ and multiple sclerosis ^144^. In both analyses P-values were corrected within each trait using the Benjamini–Hochberg procedure, and associations with q < 0.05 were considered significant.

### TF and target gene changes in cryptozoospermia

To address the question of whether a change in expression of a TF is associated with changes in expression of its target genes, TFs with a significant (p <0.05) change in expression between normal and cryptozoospermia samples and an absolute log2 fold change in expression of at least 0.5 in at least one cluster were selected. For each TF, its SCENIC+ targets were identified. For each cluster, the mean log2 fold change in expression of targets was calculated. As several clusters show an overall skew in change of gene expression, the mean log2 fold change of all genes in the given cluster was calculated and subtracted from the average change in expression of the target genes. To calculate the significance of the association in direction of change of expression of the TF and target genes, 2x2 contingency tables were calculated per TF and per cluster, for the classes “target/non-target” and “same direction/other direction”. Fisher’s exact test p-values were calculated and adjusted for multiple testing using the Benjamini-Hochberg method.

## Acknowledgements

We thank Jörg Gromoll, James Turner, and Will Scott for advice and comments on the project, and Paul-Georg Majev for advice on data analysis. We also thank Nicole Terwort for assistance with processing of testicular tissues. We thank Heidi Kersebom and Elke Kößer for histological evaluation of testicular tissues and Sabine Forsthoff for excellent support in endocrinological measurements. We thank the LMS Genomics Facility for assistance with 10X Xenium data generation. Finally, we thank the technicians of the andrology laboratory for their support in analysing ejaculate parameters. This work was supported by the Deutsche Forschungsgemeinschaft (DFG) Clinical Research Unit CRU326 ‘Male Germ Cells: from Genes to Function’ (project number 329621271; FR 4385/1-1,TU 298/4-2, LA 4064/3-2, NE 2190/3-2 and VA 1456/2 to CF, FT, SL, NN, and JMV respectively). Work in the Vaquerizas laboratory is supported by the Medical Research Council, UK (award reference MC_UP_160510 to JMV), the Academy of Medical Sciences and the Department of Business, Energy and Industrial Strategy (award reference APR3∖1017 to JMV), and the DFG Priority Programme SPP2202 ‘Spatial Genome Architecture in Development and Disease’ (project number VA 1456/1-1 to JMV). MMYC is supported by the Chain Florey programme (MC_PC_22013).

## Author contributions

JB, EI-S, IBB and MP analysed data. JB and SDP produced single-nucleus multiome libraries. IBB performed IF and H&E of spatial tissue sections, produced the tubule demarcations and generated spatial transcriptomics data. SDP performed IHC analyses. MMYC performed S-LDSC analysis. J-FC and SK carried out surgery, sample and data collection, interpretation and specification of the clinical dataset. JB, EI-S, IB, SDP, MMYC, MP, NR, CF, FT, CE, SL, NN, and JMV interpreted data. CF, SL, NN, and JMV obtained funding and supervised the study. JB, EI-S and JMV wrote the first draft of the manuscript and all authors contributed to editing the manuscript.

## Competing interests

The authors declare no competing interests.

## Data availability

Single-nucleus multiome sequencing data are available at the European Genome-Phenome Archive (EGA). Upon publication, access to the data will be available via application to the Data Access Committee. Spatial transcriptomic data will be available through the EMBL-EBI’s BioImage Archive upon publication.

## Code availability

Code will be made available via Github upon publication.

---

**Figure S1:**
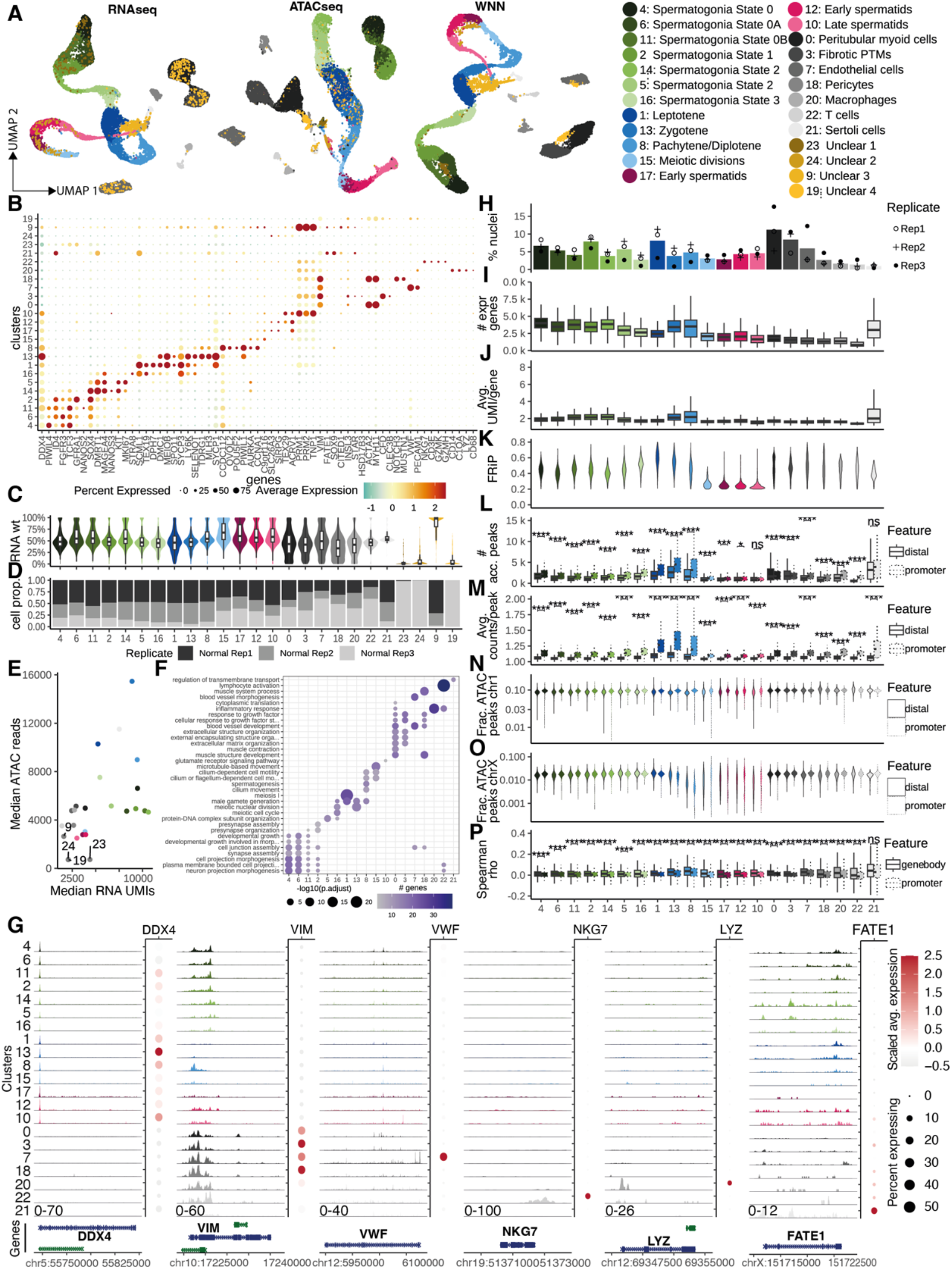
Identification and characterisation of clusters in multiome data. A UMAPs based on RNA-seq, ATAC-seq, and the weighted nearest neighbour (WNN) graph produced by combining the two modalities using Seurat. UMAPs show all clusters, including 4 “Unclear” clusters that could not be identified based on marker gene expression and were excluded from further analysis. B. Marker gene expression across clusters. Colour indicates scaled average expression, and size indicates the % of cells in the cluster with detected expression. C. (Top) RNA/ATAC-seq weight per cluster. Most clusters have a 50:50 weighting of RNA and ATAC-seq; the ‘Unclear’ clusters are skewed towards 100% weighting of one modality. D. Distribution of nuclei from each normal-sample replicate across cell types. “Unclear” clusters are enriched for nuclei from specific replicates. Specifically, cluster 23, 24 and 19 arise only from Normal 3 and Cluster 9 only from Normal 1 and 2. E. Median RNA UMIs vs Median ATAC-seq reads per cluster indicating low counts of both in the 4 “Unclear” clusters. F. Gene Ontology (Biological Process) terms that are significantly enriched in marker genes for each cluster shown in Fig.1. The topmost significant term per cluster was selected for inclusion in the plot. Size represents -log10 of the BH-adjusted p-value for enrichment of the term in the given gene set, and colour represents the number of DE genes associated with that term. Absent dots indicate no significant enrichment of that term in the given gene set. G. Each sub-panel shows cluster-specific pseudobulk chromatin accessibility tracks on the left. Expression of the marker gene of interest in the cluster shown in the dot plots on the right, where colour indicates scaled average expression and size indicates the % of cells in the cluster with detected expression. Marker genes shown are *DDX4* for germline cells, *VIM* for somatic cells, *VWF* for endothelial cells, *NKG7* for T cells, *LYZ* for macrophages and *FATE1* for Sertoli cells. H. Quantification of the contribution of cell type to the total number of cells and each biological replicate to the individual clusters. I. Total number of expressed genes in each cluster. J. Average number of RNA molecules (UMIs) per gene in each cluster. K. The fraction of reads per cell which fall inside peaks drops sharply at the meiotic divisions stage and in spermatids. Cells with less than 20% of reads in peaks were filtered out during pre-processing as these are likely to be low-quality cells. L. Total number of accessible peaks, stratified into promoter peaks (overlapping ±100 bp around annotated transcript start sites; Ensembl v98) and distal peaks, in each cluster. Statistical significance between promoter and distal peaks within each cluster was assessed using two-sided *t*-tests with Benjamini–Hochberg (BH) correction; BH adjusted p-value ****: p<= 0.0001, ***: p <= 0.001; **: p<= 0.01, *: p <= 0.05, ns: p > 0.05. M. Average number of ATAC-seq reads per peak in each cluster, split into promoter and distal peaks as in K. Statistical significance between promoter and distal peaks within each cluster was assessed using two-sided t-tests with Benjamini–Hochberg (BH) correction; BH adjusted p-value ****: p<= 0.0001, ***: p <= 0.001; **: p<= 0.01, *: p <= 0.05, ns: p > 0.05. N. Fraction of ATAC-seq reads in peaks coming from chr, split into promoter and distal peaks as in K. O. Fraction of ATAC-seq reads in peaks coming from chrX compared to chr1 (M), split into promoter and distal peaks as in K. The fraction of all ATAC-seq reads in peaks coming from chrX is consistent throughout spermatogonia and in somatic cells, but drops in zygotene and pachytene/diplotene, indicative of MSCI, and appears bimodal in meiotic divisions and spermatids, indicative of haploid cells containing either the X or Y chromosome. P. Spearman correlation between gene expression and accessibility at gene body or promoter across individual cells in each cluster. Gene bodies were defined as the genomic span of the longest annotated transcript, excluding upstream promoter regions. Promoters were defined as the 2 kb region upstream of the transcription start site (TSS) of the longest annotated transcript. Statistical significance between gene body and promoter accessibility within each cluster was assessed using two-sided *t*-tests with Benjamini–Hochberg (BH) correction; BH adjusted p-value ****: p<= 0.0001, ***: p <= 0.001; **: p<= 0.01, *: p <= 0.05, ns: p > 0.05.

**Figure S2:**
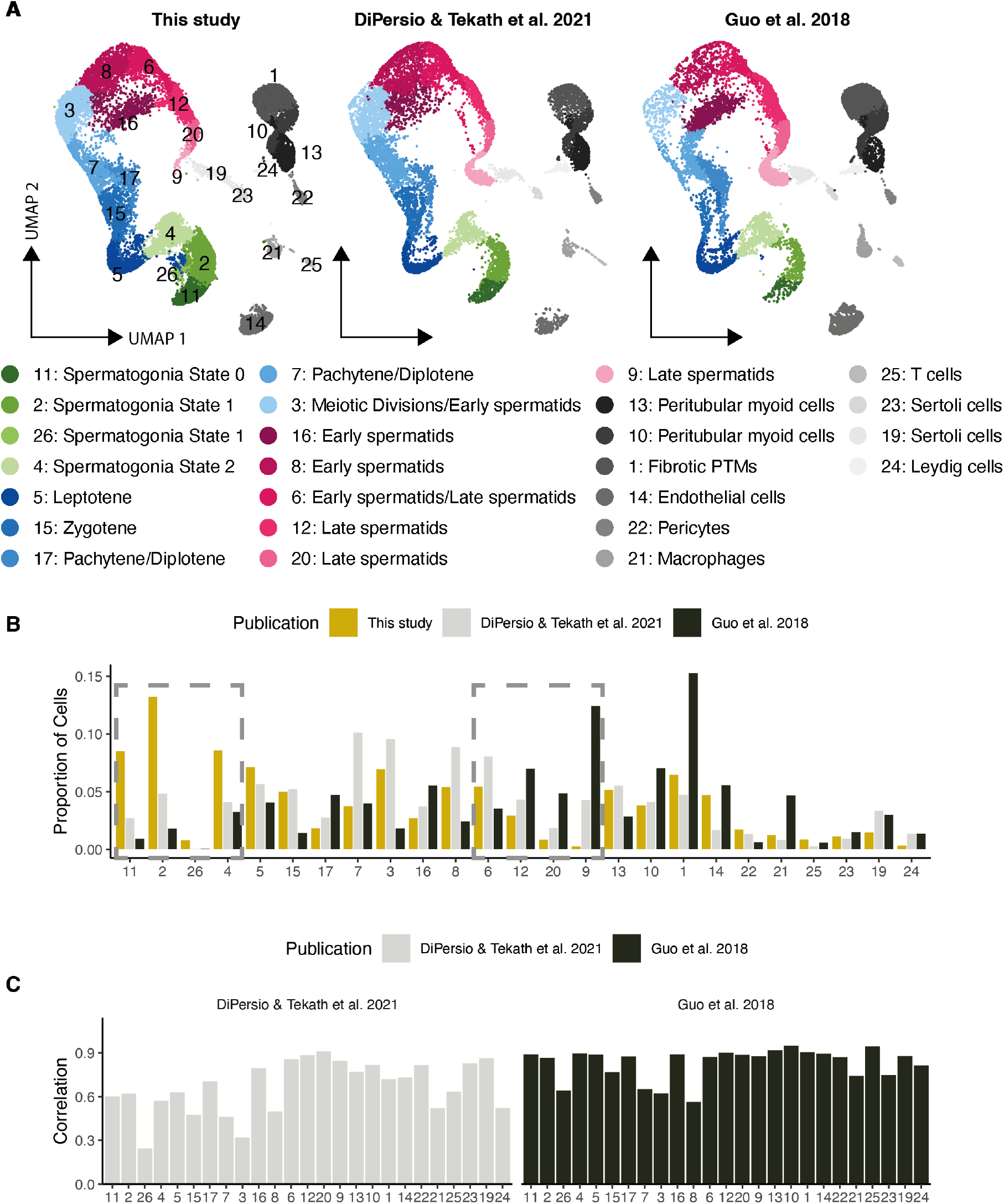
Comparison of snRNA-seq and scRNA-seq. A. UMAPs of clusters obtained from snRNA-seq (this study) and published scRNA-seq from Di Persio & Tekath et al. 2021 ^26^ and Guo et al. 2018 ^16^. B. Proportion of different cell types captured across the three datasets indicating an enrichment of spermatogonia and a depletion of spermatids in snRNA-seq compared to scRNA-seq datasets. C. Pearson correlation of gene expression between this study and Di Persio & Tekath et al. 2021 or Guo et al. 2018. All values are significant in t-test with Holm–Bonferroni adjustment with p.adj < 0.0001 for Di Persio & Tekath et al., 2021 and < 0.001 for Guo et al., 2018.

**Figure S3:**
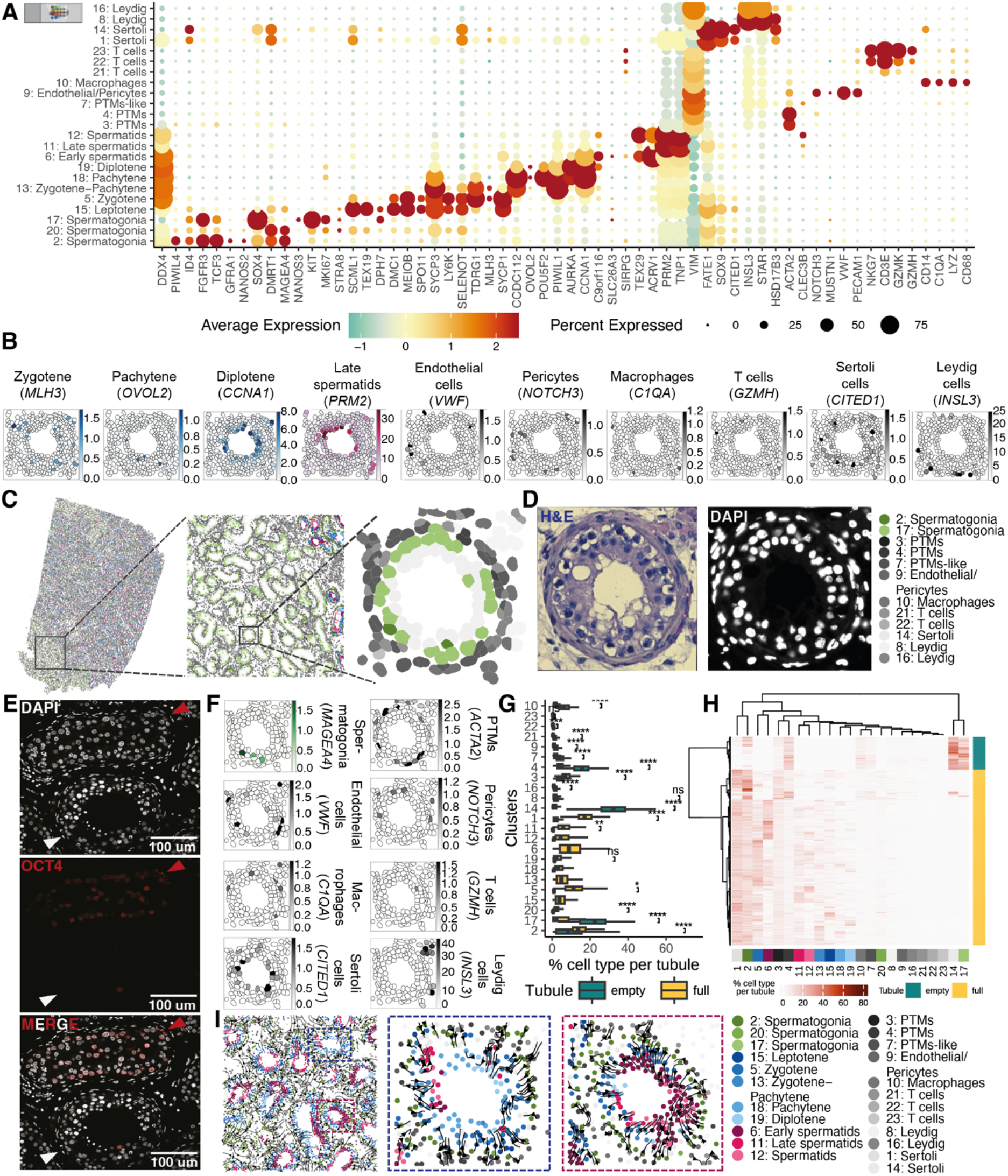
Identification and characterisation of clusters in spatial transcriptomic data from 10X Xenium In Situ platform. A. Marker gene expression across clusters in the spatial data. Colour indicates scaled average expression, and size indicates the % of cells in the cluster with detected expression. Slide icon indicates origin of data as spatial. B. Marker gene expression for the different cell types shown in the full tubule in Fig.2C. C. Scatterplot of single-cell spatial coordinates obtained from 10x Xenium In Situ platform of a human testicular tissue cross-section of 60.6 mm². Each point represents an individual cell positioned according to its spatial location within the tissue section and coloured based on celltype cluster assignment. Slide icon indicates origin of data as spatial. This is followed by zoom-in on a section of C focussing on a collection of empty tubules, followed by zoom and a further zoom-in on a single empty tubule. D. H&E staining (left) and DAPI staining (right) of the empty tubule. E. Representative immunofluorescence staining of human seminiferous tubule sections showing expression of OCT4 protein in the intratubular cells of empty tubules indicative of Germ Cell Neoplasia in Situ (GCNIS). Nuclei were counterstained with DAPI. Scale bars, 100μm. F. Marker gene expression for *MAGEA4* representing spermatogonia, *ACTA2* representing PTMs, *VWF* for endothelial cells, *NOTCH3* for pericytes, *C1QA* for macrophages, *GZMH* for T cells, *CITED1* for Sertoli cells and *INSL3* for Leydig cells in the empty tubule in C. G. Heatmap of % cell type composition per tubule classifying them into full or empty. H. % cell type composition of the different clusters per tubule for empty and full tubules. Asterisks indicate statistically significant pairwise comparisons of the same cell type between full and empty tubules; BH adjusted p-value ****: p<= 0.0001, ***: p <= 0.001; **: p<= 0.01, *: p <= 0.05, ns: p > 0.05. I. Predicted cellular trajectories inferred by SpaTrack, based on gene expression and spatial information from the spatial data and projected onto the tissue section with cell areas coloured based on cell type cluster assignment. Predicted cellular trajectories inferred by SpaTrack, within the black box highlighted in C, based on gene expression and spatial information from the spatial data and projected onto the tissue section with cell areas coloured based on cell type cluster assignment. Flow arrows indicate the inferred direction of cell-state transitions across spatial locations. H. Zoom-in of a region (highlighted in dashed blue box), showing a tubule in which spermatogenesis has progressed predominantly to the meiotic stage and a tubule in which spermatogenesis has progressed predominantly to the haploid spermatid stage (highlighted in dashed pink box).Zoom-in of a region in H (highlighted in dashed blue box), showing a tubule in which spermatogenesis has progressed predominantly to the meiotic stage and a tubule in which spermatogenesis has progressed predominantly to the haploid spermatid stage (highlighted in dashed pink box).

**Figure S4:**
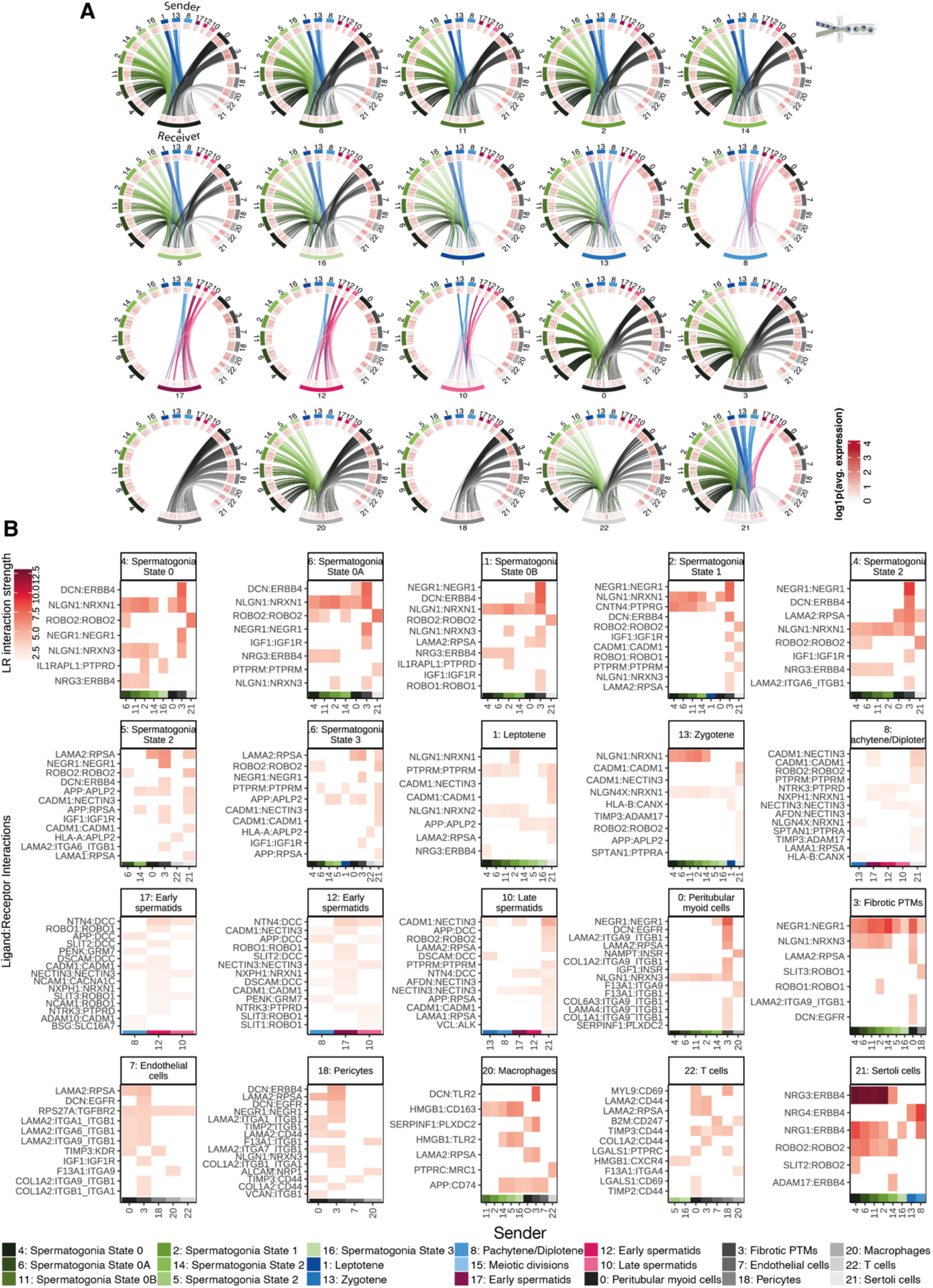
Circos plots and heatmap of LRIs in normal spermatogenesis. A. Circos plots representing significant LRIs (BH adjusted p.value < 0.01) between sender and receiver cell types inferred from the multiome data for cell type pairs with positive neighbourhood enrichment score in the spatial data. The outer-most track represents the cell type, followed by a track of heatmap of gene expression for the ligand on the sender cell types and for the receptor on the receiver cell types in the multiome data. The inner-most track represents links between the ligand receptor pairs, with the thickness of edge indicating strength of interaction and the colour indicating the sender cell type. Microfluidics icon indicates origin of data as multiome. B. Heatmap of the top 20 significant ligand–receptor interactions (LRIs), displaying interaction strength across sender cell types for each receiver cell type. The LRIs are shown on the y-axis, sender cell types are shown on the x-axis, with separate facets corresponding to receiver cell types. Microfluidics icon indicates origin of data as multiome.

**Figure S5:**
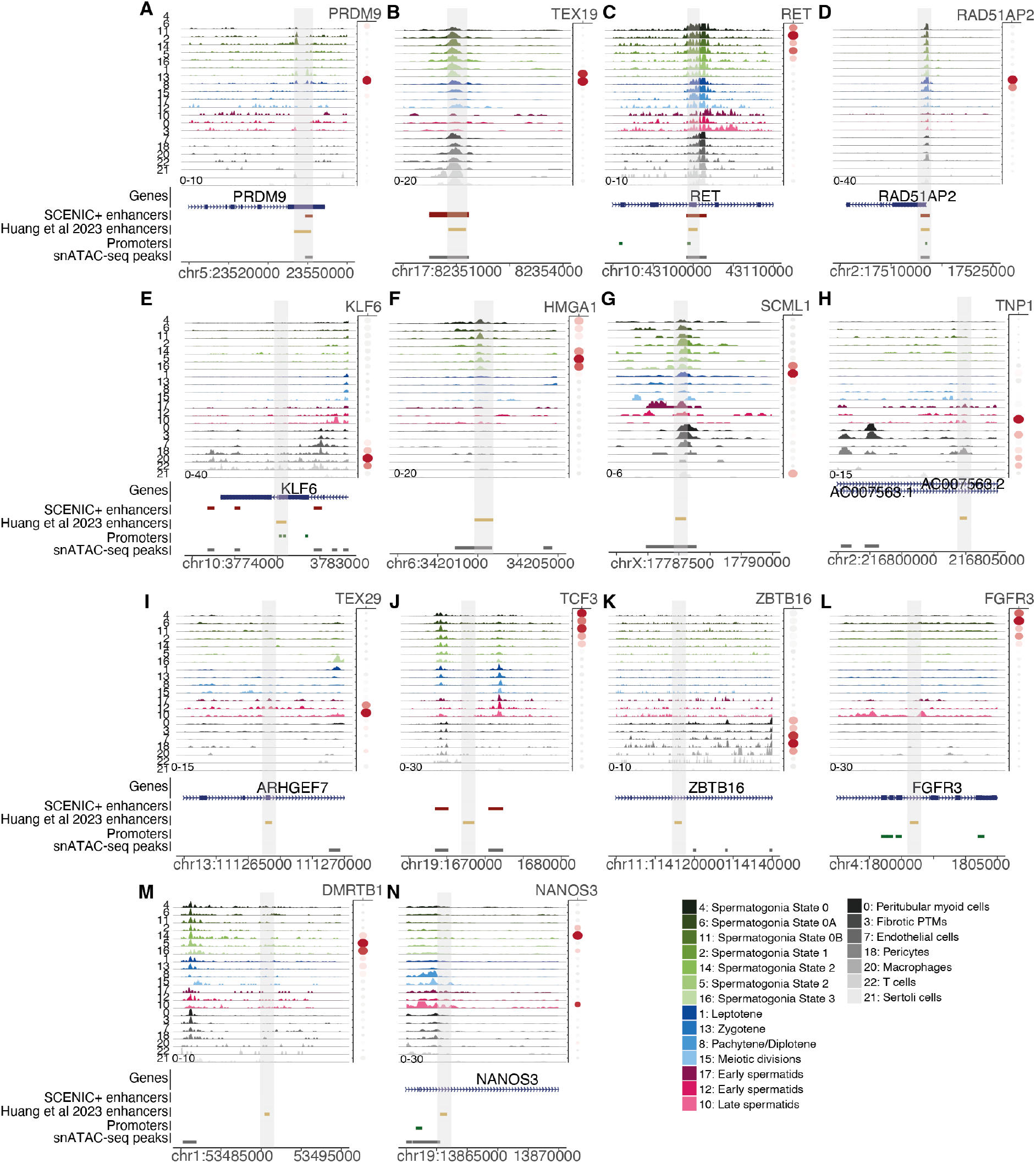
Overlap of SCENIC+ enhancers identified in this study with luciferase-activity-tested enhancers in Huang et al. 2023^25^. A-N. Each sub-panel shows cluster-specific pseudobulk chromatin accessibility tracks on the left, with gene annotations below. Regions are centred on genes of interest, which are labelled. Expression of the marker gene of interest in the cluster is shown in the dot plots on the right, where colour indicates scaled average expression and size indicate the % of cells in the cluster with detected expression. Bottom, gene annotations, predicted SCENIC+ enhancers from this study (red), luciferase-activity-tested enhancers from Huang et al. 2023 (yellow), promoters (green) and snATAC-seq peaks (grey).

**Figure S6:**
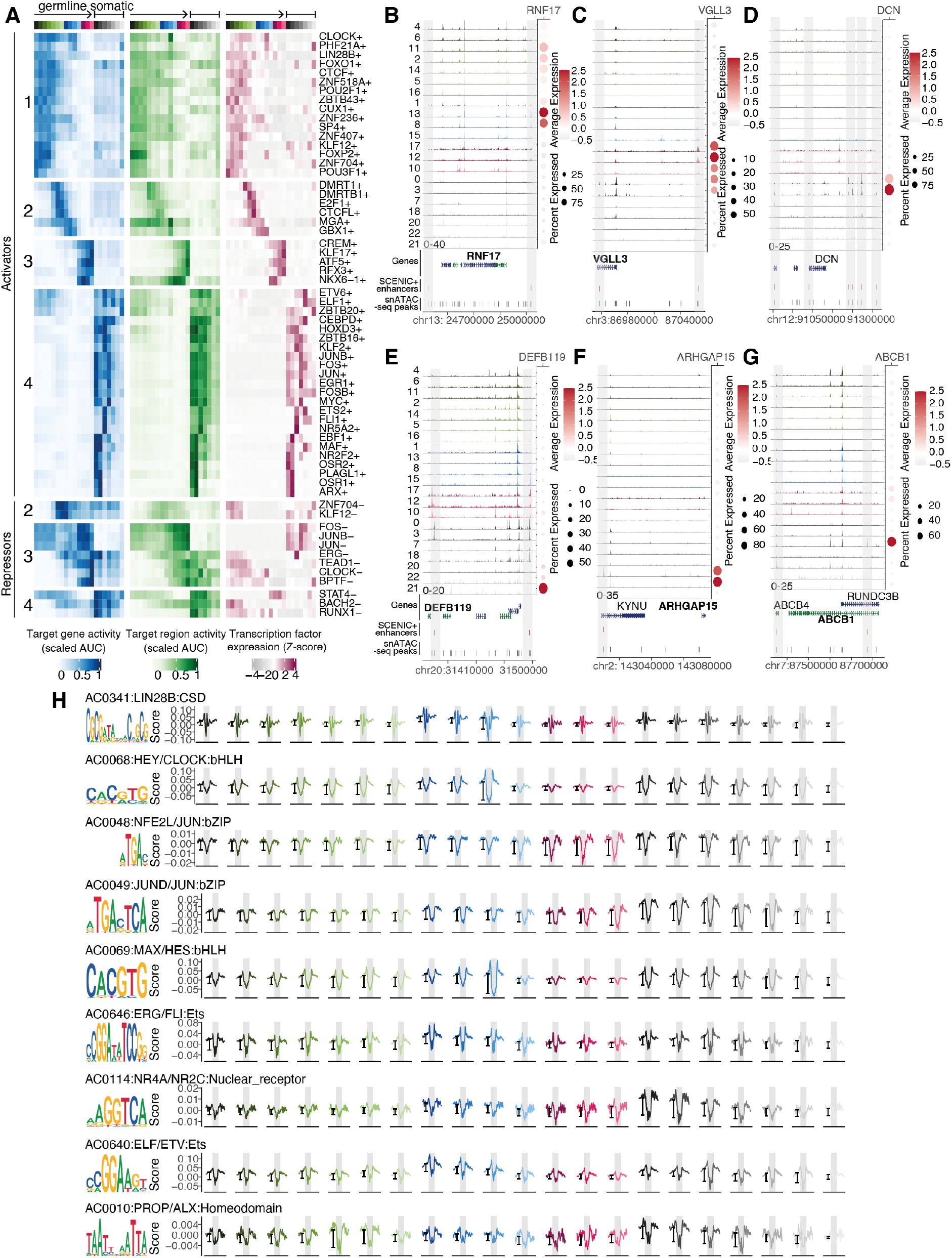
Further examples of SCENIC+ predicted enhancers and target genes in germline clusters. A. 62 high-confidence eRegulons which show dynamic activity across cell types were identified. TF name and direction of regulation are labelled on the right. Top, activator eRegulons. Bottom, repressor eRegulons. Left-to-right, panels represent eRegulon target gene expression enrichment per cluster (blue, scaled average AUC), eRegulon target region accessibility enrichment per cluster (green, scaled average AUC), and TF expression per cluster (magenta, Z-score). There are 4 distinct clusters of eRegulons active in spermatogonia (1), differentiating spermatogonia and spermatocytes (2), spermatids (3) and somatic cell types (4). B. SCENIC+ identifies two predicted enhancers for *RNF17* (red bars within grey highlights). One of the enhancers is accessible in spermatogonia, the other primarily in zygotene and pachytene/diplotene, as shown by cluster-specific pseudobulk chromatin accessibility tracks. *RNF17* expression shown as a dotplot where colour indicates scaled average expression and size indicates the % of cells in the cluster with detected expression. Bottom, gene annotations, predicted enhancers (red), and scATAC-seq peaks (grey). C. Same as B, for *VGLL3* (red bars within grey highlights). The lefthand enhancer is predicted to be regulated by EBF1 and NR2F2 and the righthand enhancer by ATF5 and CREM. D. Same as B, for *DCN* (red bars within grey highlights). All of the enhancers are accessible in PTMs and/or fibrotic PTMs, as shown by cluster-specific pseudobulk chromatin accessibility tracks. E. Same as B, for *DEFB119* (red bars within grey highlights). The enhancers are accessible in Sertoli cells, as shown by cluster-specific pseudobulk chromatin accessibility tracks. F. Same as B, for *ARHGAP15* (red bars within grey highlights). This enhancer is accessible in macrophages, as shown by cluster-specific pseudobulk chromatin accessibility tracks. G. Same as B, for *ABCB1* (red bars within grey highlights). The enhancers are accessible in endothelial cells, as shown by cluster-specific pseudobulk chromatin accessibility tracks. H. Selected motifs targeted by LIN28B (AC0341), CLOCK (AC0068), JUN (AC0048), FOS/FOSB/JUN (AC0049), MYC (AC0069), ERG (AC0646), NR2F2 (AC0114), RUNX1/ELF1 (AC0640) and ARX (AC0010). TOBIAS footprinting profiles indicate LIN28B, CLOCK, and JUN are strongly bound in pachytene/diplotene spermatocytes; FOS, FOSB, JUN, MYC, ERG, NR2F2, RUNX1, ARX are bound strongly in PTMs and fibrotic PTMs; ELF1 is bound strongly in Sertoli cells and in macrophages and ETS2 and FLI1 are bound strongly in endothelial cells.

**Figure S7:**
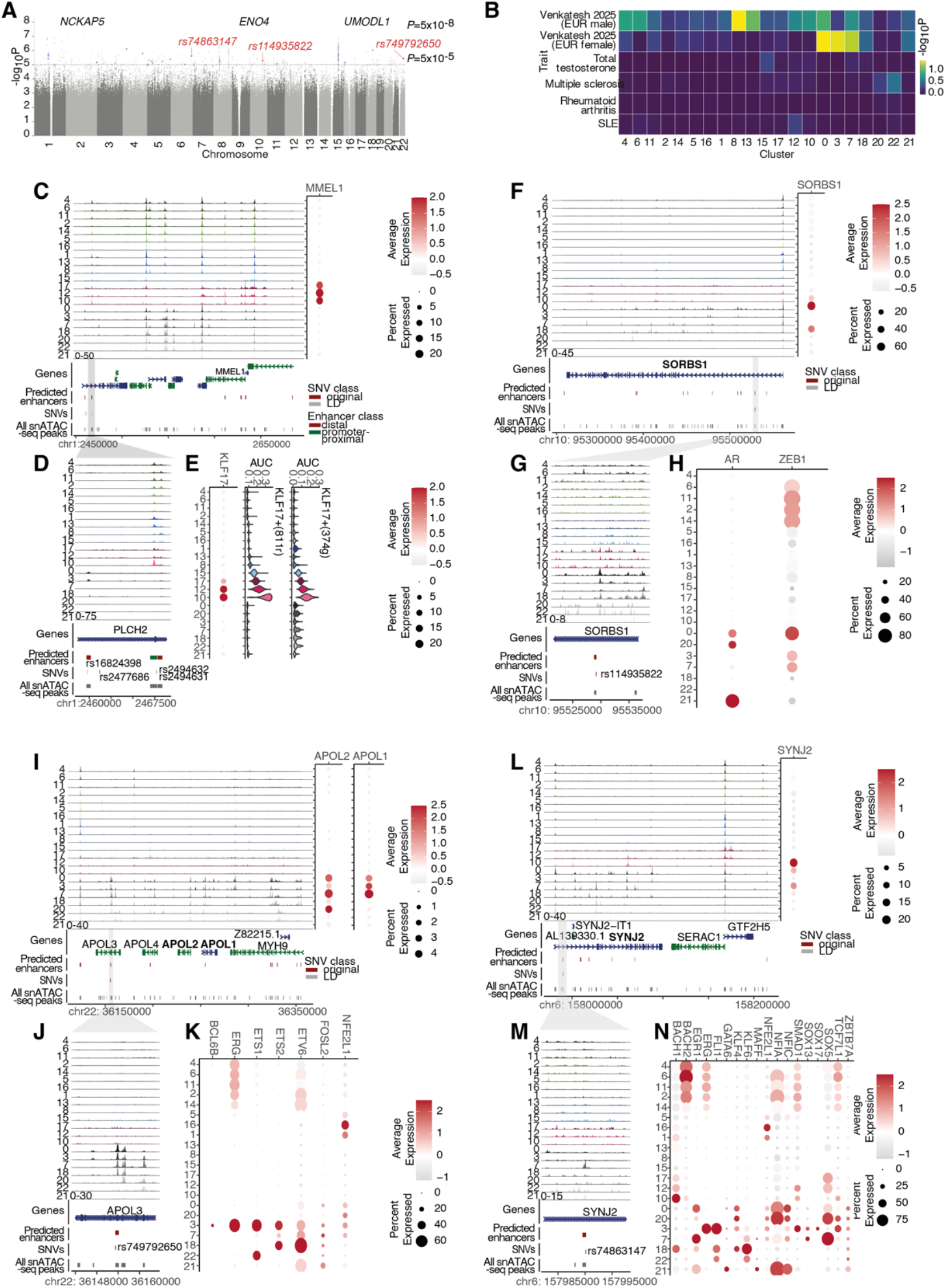
SCENIC+ results identify candidate target genes and enhancers for GWAS loci. A. Manhattan plot of the multi-ancestry male infertility meta-analysis from Venkatesh et al, 2025 ^41^ (10,886 cases and 995,982 controls; 28,589,555 biallelic autosomal variants). Summary statistics from whole-genome regression analyses were meta-analyzed using fixed-effect inverse-variance weighting in the METAL software to produce the displayed *P* values. Summary statistics for chromosome X were not publicly available. Chromosomal position (GRCh38) is shown along the *x*-axis; association strength is plotted on the *y*-axis as −log₁₀(P). Each point represents a single variant. The dashed line denotes the genome-wide significance threshold (*P* = 5 × 10⁻⁸) with the three significant loci annotated by nearest gene. The dotted line indicates suggestive association (*P* < 10⁻⁵). Variants highlighted in red overlap with SCENIC+ enhancers. Variants highlighted in blue overlap testis-specific ATAC-seq peaks. B. Cell-type-specific partitioned SNP heritability across European male (6,885 cases and 410,578 controls) and female (40,024 cases and 665,658 controls) infertility and other selected traits associated with male reproductive health. Stratified LD score regression coefficient *P*-values for 21 cell types (columns) across six traits (rows). Cell-type annotations were MACS2 peak calls from snATAC-seq generated from adult human testes, each analysed conditional on the baseline model (v2.2). Tile fill shows −log_10_(P) for the cell-type-specific regression coefficient τc, a one-sided test of whether the annotation contributes to trait heritability beyond the baseline annotations. Colour is capped at the 99^th^ percentile of the pooled −log_10_(P) distribution; tiles at the ceiling are shown in the terminal colour. GWAS summary statistics shown for male and female infertility ^41^, total testosterone ^141^, systemic lupus erythematosus ^142^, rheumatoid arthritis ^143^ and multiple sclerosis ^144^.). No annotation was significant after correction for multiple testing (false discovery rate q < 0.05, applied within each trait across all 21 cell types). C. *MMEL1* locus. Left, cluster-specific pseudobulk chromatin accessibility tracks. Right, *MMEL1* expression per cluster. Bottom, gene annotations, SCENIC+ predicted distal enhancers for *MMEL1*, SNVs, and snATAC-seq peaks. Predicted enhancers that satisfied the non-promoter-overlapping criteria are shown in dark red, whereas promoter-overlapping predicted enhancers excluded from the final SCENIC+ enhancer set are shown in green. The lead SNV (dark red) is rs2477686 identified as associated with male infertility by Hu et al. 2012 ^59^; variants in high LD (R^2^ > 0.8) with this SNV are shown in grey. D. Top, close up of part of the region in A. Two variants in high LD (R^2^ > 0.8) with rs2477686 (rs2494631 and rs2494632), overlap an snATAC-seq peak identified by SCENIC+ as a putative enhancer of *MMEL1*, predicted to be regulated by KLF17, but excluded from the final enhancer set because it overlaps a promoter (green). A variant in high LD (R^2^ > 0.8) with rs2477686 (rs16824398), overlaps a distal enhancer predicted to regulate both *MMEL1* and *TNFRS14*, as described in Fig. 4. E. KLF17 eRegulon activity across single nuclei, as measured by target gene expression enrichment (top) and target region chromatin accessibility enrichment (bottom). *KLF17* expression across clusters; colour indicates scaled average expression, and size indicates the % of cells in the cluster with detected expression. F-H. Same as C-E, for *SORBS1*. I-K. Same as C-E, for *APOL2* and *APOL1*. L-N. Same as C-E, for *SYNJ2*.

**Figure S8:**
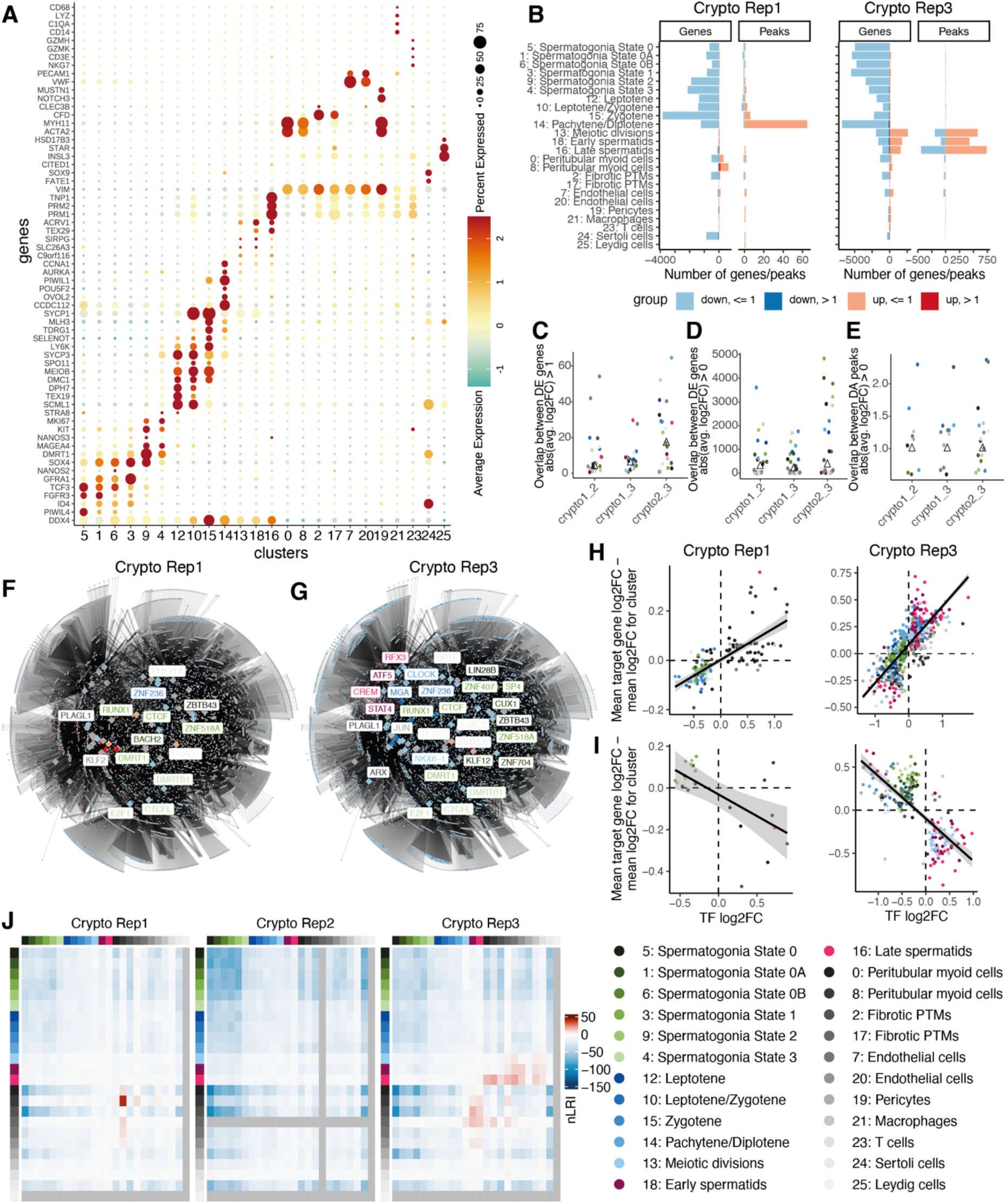
Identification and characterisation of clusters in the combined data set (normal + cryptozoospermia). A. Marker gene expression across clusters. Colour indicates scaled average expression, and size indicates the % of cells in the cluster with detected expression. B. Numbers of differentially expressed genes (left) and differentially accessible peaks (right) in each cluster, comparing normal spermatogenesis and Crypto Rep1 or Crypto Rep3. C. Overlap of DE genes identified in Crypto Rep1 vs Normal and Crypto Rep2 vs Normal comparisons or Crypto Rep1 vs Normal and Crypto Rep3 vs Normal or Crypto Rep2 vs Normal and Crypto Rep3 vs Normal. DE genes were defined as those with abs(log2FC) > 1. D. Same as C, showing overlap of DE genes or (E.) DA peaks. DE genes/DA peaks were defined as those with abs(log2FC) > 0. F-G. Differentially expressed genes between normal spermatogenesis and Crypto Rep1 or Crypto Rep3 shown in the context of the high-confidence gene regulatory network (Fig. 3D). TFs are shown as large diamonds and all other genes as smaller circles. Genes that are significantly differentially expressed in any cluster are shown in red and all other genes in grey. Edges represent a regulatory relationship between a TF and a target gene inferred by SCENIC+. Differentially expressed TFs are labelled and coloured according to the cluster in which they have highest expression in the normal spermatogenesis samples. H. Activator TFs significantly differentially expressed in any cluster between normal and Crypto Rep1 (left) or Crypto Rep3 (right) (p < 0.05) are shown. Each point represents a TF and its targets in a given cluster; the logFC of the TF is represented on the x axis and the difference in mean log2FC between target genes and all genes is plotted on the y axis, as explained in Fig.5E. Clusters are coloured according to the key in panel D. Points with a Fisher test p-value of < 0.05 for the association between direction of change in expression for the TF and direction of change in expression of its targets are shown. I. Same as E, for repressor TFs. J. Heatmap of number of differential LRIs between Crypto and Normal spermtogenesis. Rows and columns are arranged in developmental order with the germ cells followed by the somatic cells. Taking number of significant differential interactions between Crypto and Normal samples (BH padj < 0.01). The Normal samples contained fewer than 20 cells in cluster 25: Leydig cells and Crypto Rep2 contained fewer than 20 cells in cluster 17: Fibrotic PTMs, Therefore, these cell types were excluded from the differential LRI analysis, and the corresponding rows and columns are shown in grey.

**Figure S9:**
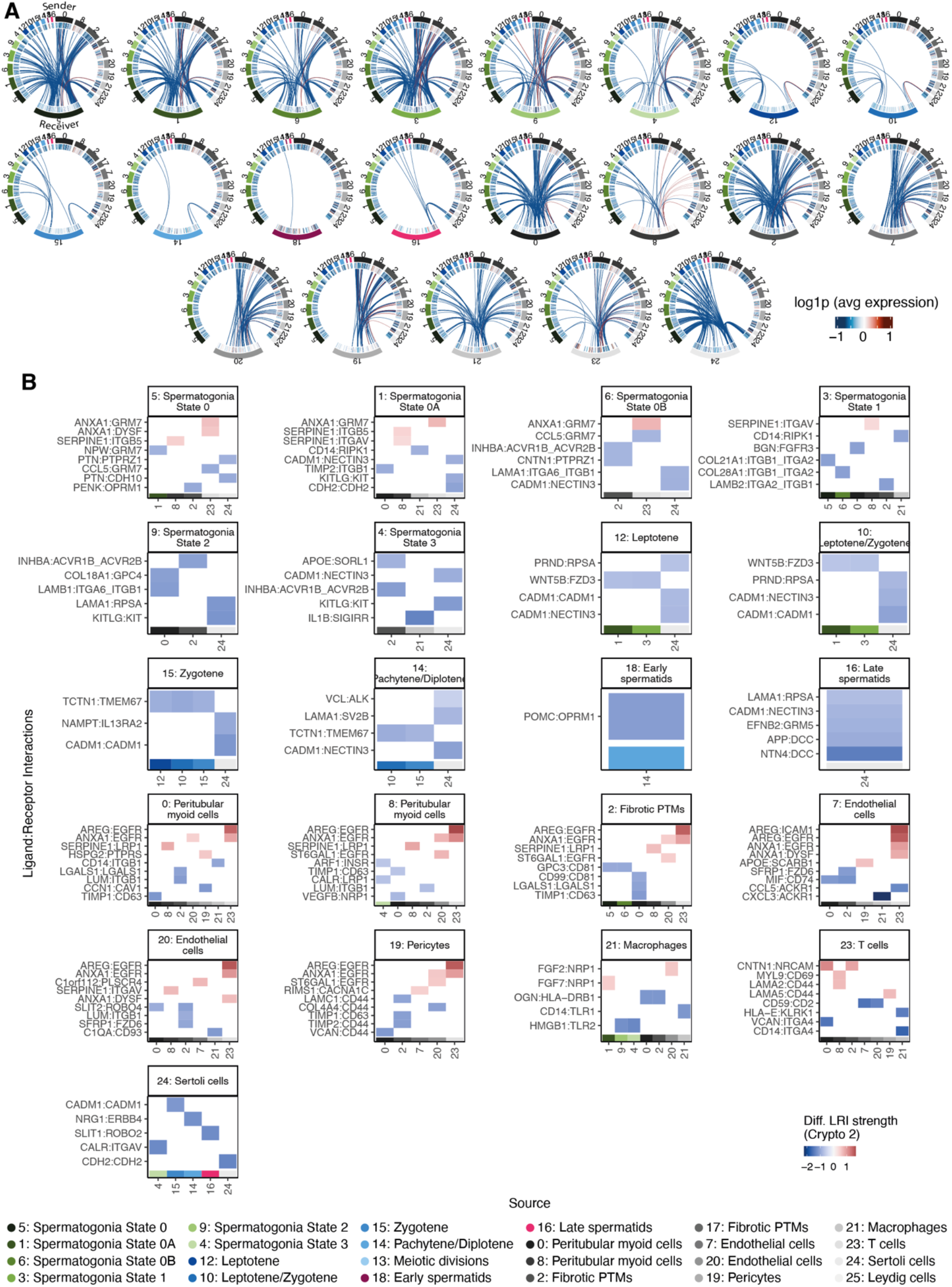
Circos plots and heatmap of differential strength of LRIs in Crypto Rep2 vs normal spermatogenesis. A. Circos plots representing significantly differential LRIs (BH adjusted p.value < 0.01) between sender and receiver cell types inferred from the multiome data for cell type pairs with positive neighbourhood enrichment score in the spatial data. The outer-most track represents the cell type, followed by a track of heatmap of log2FC of gene expression between crypto and healthy conditions for the ligand on the sender cell types and for the receptor on the receiver cell types in the multiome data. The inner-most track represents links between the ligand receptor pairs, with the thickness of edge indicating log2FC of strength of interaction between crypto and healthy conditions and the colour indicating upregulated (red) or downregulated (blue) LRIs between crypto and healthy conditions. B. Heatmap of the top 5 up- and downregulated ligand–receptor interactions (LRIs) between Crypto Rep2 and normal conditions, across sender cell types for each receiver cell type. The LRIs are shown on the y-axis, sender cell types are shown on the x-axis, with separate facets corresponding to receiver cell types. Ligands and receptors discussed in the text are indicated in bold.

## Notes

### Competing Interest Statement

The authors have declared no competing interest.

